# Nanobodies against *Plasmodium vivax* AMA1 block parasite invasion

**DOI:** 10.64898/2026.11.01.763489

**Authors:** Jill Chmielewski, Li-Jin Chan, Li Lynn Tan, Jaison D Sa, Amy Adair, Jerry Jiaheng Chen, Chunyi Qian, Phillip Pymm, Frankie M. T. Lyons, Kathleen Zeglinski, James G. Beeson, Christopher L. King, Wai-Hong Tham, Melanie H. Dietrich

**Affiliations:** Walter and Eliza Hall Institute of Medical Research, Parkville, Victoria, Australia; Department of Medical Biology, The University of Melbourne, Parkville, Victoria, Australia; Burnet Institute, Melbourne, Victoria, Australia; Department of Medicine and Department of Infectious Diseases, The University of Melbourne, Parkville, Victoria, Australia; School of Translational Medicine, Monash University, Melbourne, Victoria, Australia; Centre for Global Health and Diseases, Case Western Reserve University, Cleveland, Ohio, United States of America; Veteran Affairs Research Service, VA Medical Center, Cleveland, Ohio, United States of America; Research School of Biology, The Australian National University, Canberra, Australian Capital Territory, Australia

**Keywords:** malaria, Plasmodium vivax, parasite invasion, crystallography, nanobody

## Abstract

*Plasmodium vivax* is the most widespread malaria species to infect human populations. *P. vivax* invades young red cells called reticulocytes to replicate and proliferate. An essential protein in parasite invasion is the Apical Membrane Antigen 1, PvAMA1, which forms the tight junction between parasite and red cell membrane through its interaction with Rhoptry Neck Protein 2, PvRON2. Here we report the identification of PvAMA1 nanobodies using phage display in combination with Sanger sequencing, next-generation sequencing and structure prediction tools. We show that these nanobodies bind with high affinities and across several different epitopes on PvAMA1. Several nanobodies recognize PvAMA1 in merozoites of late-stage schizonts and inhibit parasite invasion of red blood cells. These growth inhibitory nanobodies bind to conformational epitopes of PvAMA1 and most bind to a site distinct from a known potent human monoclonal antibody 826827. Crystal structures of two inhibitory nanobodies in complex with PvAMA1 show that they bind novel neutralizing epitopes which are outside the hydrophobic groove that contacts PvRON2. Furthermore, we show that two other PvAMA1 nanobodies also inhibit *Plasmodium falciparum* invasion. These results demonstrate the potential of PvAMA1 nanobodies to inhibit malaria parasite invasion across multiple *Plasmodium* species.

**Author Summary:** Malaria parasites are exquisitely adapted for invasion and survival within human red blood cells. Merozoites are the form of the malaria parasites that invade red cells. Parasite invasion begins with initial attachment of the merozoite to the red blood cell surface, followed by tight junction formation between the parasite and the red blood cell membrane. Active invasion proceeds as the tight junction moves from the apical to posterior pole of the merozoite and pulls the parasite into the red blood cell. In *Plasmodium vivax*, PvAMA1 is critical for tight junction formation during parasite invasion. Here we describe the identification of nanobodies that bind to PvAMA1 at different sites using phage display combined with Sanger sequencing, next-generation sequencing and structure prediction approaches. We show that PvAMA1 nanobodies potently inhibit malaria parasite invasion in multiple *Plasmodium* species, demonstrating their potential to reduce malaria infection of red blood cells.

## Introduction

*Plasmodium vivax* is one of the most widespread human malaria parasite species with around 3 billion people at risk. *P. vivax* preferentially invades reticulocytes in a multi-step process [1–4]. A merozoite makes initial contact with the red blood cell surface and then reorientates to position its apical pole with the red blood cell membrane to interact with red blood cell receptors. Several red cell receptors have been identified to be involved in *P. vivax* invasion, which include the Duffy antigen receptor for chemokines (DARC), CD98 and Transferrin Receptor 1 (TfR1) [5–7]. Subsequently, a tight junction is formed between the merozoite surface and red blood cell membrane and the parasite’s actomyosin motor actively pulls the merozoite into the red cell, completing the invasion process [8–15].

Key proteins in tight junction formation are Apical Membrane Antigen 1 (AMA1) and its interacting partner Rhoptry Neck Protein 2 (RON2). Both these proteins have also been found in *Toxoplasma*, *Babesia* and *Neospora* [16,17]. Since there is no long-term *in vitro* culture for *P. vivax*, most of its invasion process is inferred from what is known in other *Plasmodium* species. In *Plasmodium* parasites, AMA1 is expressed in the micronemes and around the time of schizont rupture and merozoite release, it is translocated to the parasite’s membrane [18,19]. Immunofluorescence assays using AMA1 antibodies show mostly apical localization, however, it can be detected circumferentially on the merozoite surface [18–20]. RON2 is part of the RON complex which, in *Plasmodium*, also contains RON4 and 5 [21,22]. At the start of the malaria parasite invasion, the RON complex is secreted from the rhoptries and RON2 is embedded in the red blood cell membrane [16]. RON2 forms a complex with RON4 and 5, which localize to the red blood cell cytosol, forming a positively charged platform which closely associates with the red blood cell membrane [23,24]. RON2 and AMA1 interact to initiate formation of the tight junction.

Antibodies to AMA1 have been shown to inhibit invasion in *P. falciparum*, *P. vivax*, and other *Plasmodium* species [25–30] further highlighting the importance of AMA1 in red blood cell invasion. AMA1 has been a focus of vaccine development for *P. falciparum* and *P. vivax* [31,32], and has also been considered as a target for therapeutic monoclonal antibodies and small molecule inhibitors [27,33].

AMA1 is composed of a signal sequence, three extracellular domains (D1 to D3), a single transmembrane region and a cytoplasmic domain. A short extra-cellular loop of RON2 binds to the hydrophobic groove in AMA1 D1, which is fully exposed when a mobile loop of AMA1 D2 is displaced [25,34,35]. The AMA1 D2-loop also has an important role in stabilizing the AMA1-RON2 complex by making direct contact with the red blood cell membrane [24,36]. Since the hydrophobic groove in AMA1 is important for RON2 binding, this region has been a target of inhibitory monoclonal antibodies against *Plasmodium* AMA1 [25,26,29,37].

While there are numerous monoclonal antibodies described against PfAMA1, only a handful of monoclonal antibodies that recognize PvAMA1 have been described using structural biology approaches. A human monoclonal antibody 826827, which was isolated from a Cambodian individual with prior infection, blocked *P. vivax* invasion of reticulocytes in several clinical isolates as well as invasion of sporozoites into liver cells [27]. The crystal structure of 826827 with PvAMA1 showed that the binding footprint of the antibody partially overlapped with the hydrophobic groove where RON2 binds [27]. WD34 is an i-body, which is a humanized single-domain antibody-like molecule, that was isolated against *P. falciparum* AMA1 (PfAMA1) using phage display [28]. It was also shown to bind to PvAMA1 with an affinity of 119 nM, which was approximately 30-fold lower than its binding to PfAMA1. Since there is no long-term *in vitro* culture of *P. vivax,* invasion inhibition was tested using a *P. falciparum* transgenic line engineered to express PvAMA1 (called W2-PvAMA1) [28,38]. WD34 could inhibit invasion of these transgenic *P. falciparum* parasites expressing PvAMA1 with an IC_50_ of 128 μg/mL [28]. Crystal structures of WD34 with PfAMA1 or PvAMA1 show that the i-body binds across the conserved hydrophobic groove in AMA1 [28]. Antibodies F8.12.19 and F8.1.1 were raised against PvAMA1 and both bind a similar epitope in D3 [39,40]. Antibody F8.12.19 was shown to have cross-species specificity as it detected AMA1 in *P. falciparum, P. cynomolgi, P. berghei* and *P. knowlesi*. However, it was non-inhibitory in growth inhibition assays performed with *ex vivo P. cynomolgi* infected red blood cells taken from infected rhesus monkeys [39].

To identify more monoclonal antibodies against PvAMA1 that have potential therapeutic use, we characterized nanobodies against PvAMA1 and showed they bound with high affinity and blocked parasite invasion. Structural analyses of two inhibitory nanobodies show that they bind to a site that is distinct from the hydrophobic groove on PvAMA1. In addition, we show that several PvAMA1 nanobodies also block *P. falciparum* parasite invasion.

## Results

### Isolation of nanobodies against PvAMA1 using phage display

To identify nanobodies against PvAMA1, we immunized an alpaca with recombinant PvAMA1 (Palo Alto variant) and generated the associated nanobody phage library. After two rounds of phage display, we observed 93 of 94 phage supernatants bound to PvAMA1 by ELISA (S1A Fig). Sanger sequencing of these positive hits identified 14 unique nanobodies (2688 – 2701), which were representatives from clonal groups determined based on a complementarity determining region 3 (CDR3) sequence similarity of ≥ 85% (Fig 1A and S1 Data). The length of the CDR3 sequences ranged between 17 and 21 residues (Fig 1A and S1 Data). Using bio-layer interferometry (BLI), we determined the binding affinities of the nanobodies to be between *K*_D_ 1.71 and 113.50 nM (Fig 1B, S1B Fig and S1 Data). Epitope binning showed that all 14 nanobodies bound the same epitope bin, which we will refer to as Group A (S1C Fig). Using a selection of three nanobodies as representatives, we observed that 2691, 2693 and 2700 bound to a different site to 826827, a previously characterized inhibitory PvAMA1 human monoclonal antibody which has a binding footprint that partially overlaps with the hydrophobic groove where RON2 binds (Fig 1C) [27]. As expected, the negative control SARS-CoV-2 nanobody WNb27 did not bind to PvAMA1 (Fig 1C and S1B Fig) [41].

**Figure 1.**
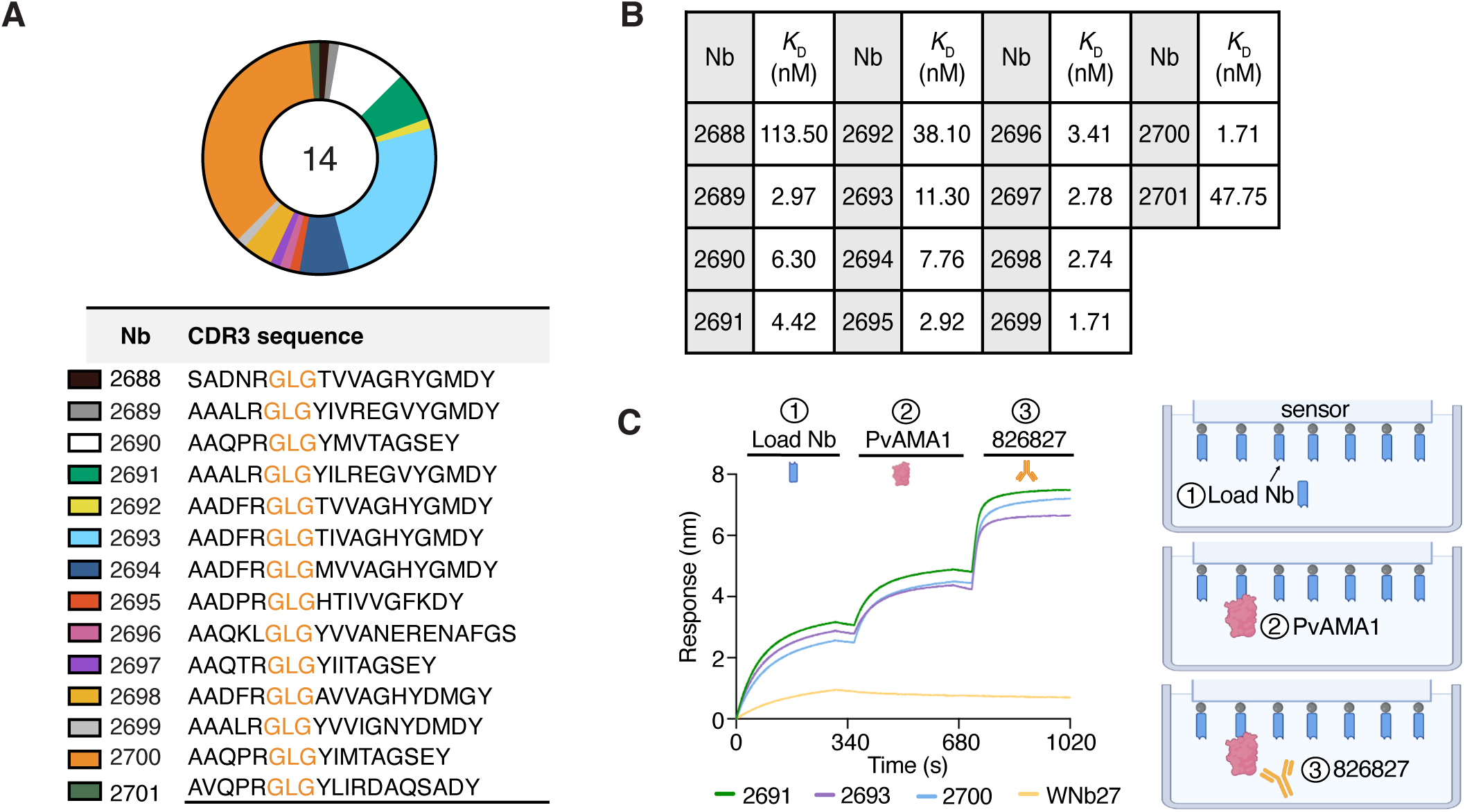
PvAMA1 nanobodies identified via Sanger sequencing. **(A)** Overview of the 14 PvAMA1 clonal groups identified by phage display and Sanger sequencing (above). The representative nanobodies selected for each clonal group and their CDR3 sequence (below). **(B)** Binding affinities (*K*_D_) of Sanger sequencing selected nanobodies to PvAMA1 determined by bio-layer interferometry (BLI). Values represent means from two independent experiments (n=2). **(C)** Epitope binning experiment using BLI with nanobodies (2691, 2693, 2700), PvAMA1 Palo Alto and 826827. His-tagged nanobodies (blue) immobilized to NTA biosensor were loaded at 5 µg/mL shown as an increased signal (nm) from 0-300 s (Step 1). Nanobody bound sensors were dipped into PvAMA1 Palo Alto (pink) at 100 nM until saturation was achieved as indicated by the second positive binding response from 360-660 s. Nanobody-loaded sensors bound with PvAMA1 Palo Alto were dipped in 100 nM of antibody 826827 (orange) from 700-1000 s to determine binding in the presence of nanobody (Step 3). Non-competitive binding is indicated as a positive binding signal in the 700-1000 s portion of the plotted curves. Nanobody measurements were plotted in solid lines: 2691(green), 2693 (purple), 2700 (blue) and WNb27 (yellow). WNb27 a SARS-CoV-2 nanobody was used as a negative control. Graphical representation of experimental setup is depicted on the right.

### Identification of nanobodies against PvAMA1 using next-generation sequencing and AI-assisted selection

To identify more nanobodies against PvAMA1 that bind to different sites from Group A nanobodies, we used next-generation sequencing (NGS) and structure prediction analyses. NGS was used to identify nanobody sequences present across the different rounds of phage display, and these sequences were clustered based on > 90% similarity in their CDR3 using Alpseq (Fig 2A) [42]. In the top 100 most abundant CDR3 sequences from NGS, we observed the presence of a GLG motif in 56 of the unique CDR3s from round 1 and 85 of the unique CDR3s from round 2 of phage display selection (S2 Data). Interestingly, this GLG motif was also present in the initial 14 nanobodies identified via Sanger sequencing (Fig 1A and S1 Data). Given the strong enrichment of this motif across our datasets, GLG-containing nanobodies were excluded from subsequent structure prediction analyses to avoid over-representation of this sequence class and to increase the diversity of candidates assessed. In the top 100 most abundant CDR3 sequences of round 1, we identified two CDR3 sequences with incomplete sequence coverage of the full length nanobody sequence. As such, these nanobodies were excluded from subsequent structure prediction analyses (S2 Data). We then used structure prediction methods to analyze 42 nanobodies identified in round 1 for their potential to engage the highly conserved hydrophobic groove in PvAMA1 that binds RON2.

**Figure 2.**
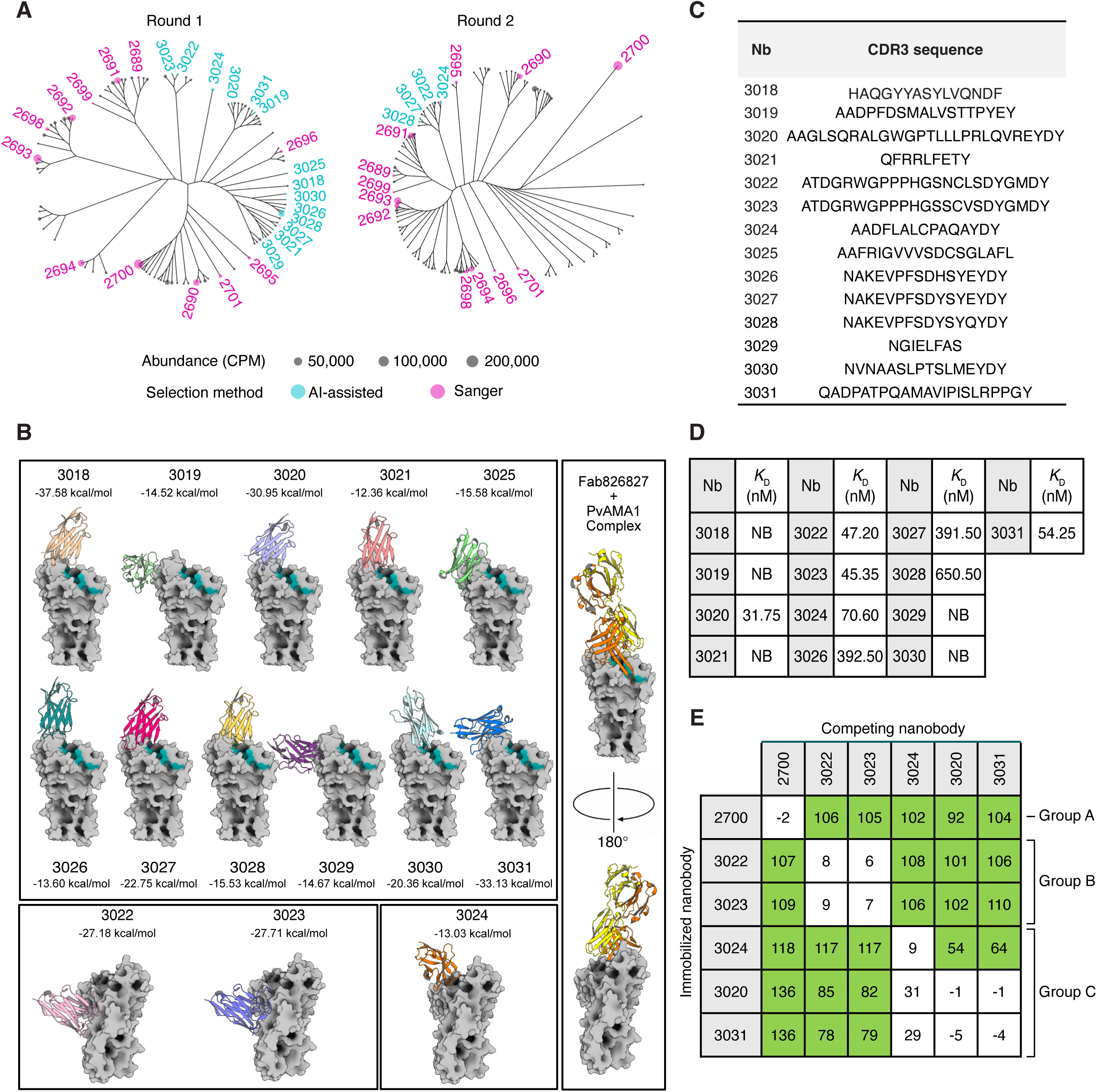
AI-assisted selection of PvAMA1 nanobodies from NGS analysis. **(A)** Cladograms of the 100 most abundant nanobodies from Next-Generation Sequencing (NGS) of round 1 and round 2 phage display libraries after biopanning. The PvAMA1 nanobodies identified by Sanger sequencing are labelled in pink, and those selected using structure prediction methods are labelled in teal. Tree tips are scaled relative to abundance by counts per million (CPM). **(B)** AlphaFold 3 predictions of nanobody–PvAMA1 complexes. The highest-confidence predicted model for each of the 14 selected AI nanobodies is shown in complex with PvAMA1, with structures grouped according to their predicted binding sites. Residues forming the hydrophobic groove where RON2 binds are highlighted in teal. The corresponding predicted FoldX interaction energy is shown beneath each nanobody. The published Fab 826827–PvAMA1 complex structure (PDB ID 9DX6) is included for reference. **(C)** CDR3 sequences of the 14 AI-selected nanobodies. **(D)** Binding affinities (*K*_D_) of AI-selected nanobodies to PvAMA1 determined by BLI. Values represent means from two independent experiments (n=2). NB, no binding. **(E)** Epitope binning experiments of AI-selected NGS nanobodies and 2700 using BLI with PvAMA1 Palo Alto. 2700 was included as a representative from the Group A nanobodies. Immobilized nanobodies are indicated on the left column and nanobodies pre-incubated with PvAMA1 Palo Alto at a 6:1 molar ratio are indicated on the top row. Binding of PvAMA1 pre-mixed with nanobody was calculated relative to PvAMA1 binding alone, which was assigned to 100%. The green and white boxes represent non-competing and competing nanobodies, respectively. Epitope bins are indicated on the right as Group A, B and C accordingly.

AlphaFold 3 modelling [43] predicted 19 out of the 42 nanobodies to bind within or adjacent to the hydrophobic groove where RON2 binds. From these, we manually selected 11 candidates that produced plausible docking poses with favorable predicted interaction energies (Δ*G* < -10 *kcal/mol*) using FoldX [44] (Fig 2B, 2C and S3 Data). Among the selected candidates, 3026, 3027 and 3028 fell within a single clonal group. In addition, we selected 3024 as its predicted binding epitope lay adjacent to the hydrophobic groove but was spatially distinct from the binding sites predicted for the other candidates. Finally, 3022 and 3023 were selected as they were predicted to bind distal from the hydrophobic groove (Fig 2B, 2C). In total, this process yielded 14 candidates (3018 – 3031) for experimental validation (Fig 2B, 2C and S3 Data). We were able to express all nanobodies except for 3025, which was not carried forward for further analyses.

Using BLI, we observed that five of these nanobodies did not bind PvAMA1, namely 3018, 3019, 3021, 3029 and 3030 (Fig 2D, S1 Data and S2 Fig). Nanobodies 3026, 3027 and 3028 bound PvAMA1 with low affinity, with *K*_D_ values between 391.50 and 650.50 nM (Fig 2D, S1 Data and S2 Fig). For the other five nanobodies (3020, 3022, 3023, 3024, and 3031), we determined higher binding affinities with *K*_D_ values between 31.75 and 70.60 nM (Fig 2D, S1 Data and S2 Fig). We performed epitope binning experiments with the five higher affinity nanobodies and 2700 as a representative from the Group A nanobodies identified by Sanger sequencing (Fig 2E). We did not perform the epitope binning for the lower affinity nanobodies as the binding signal was too low at the amount of PvAMA1 used, making it difficult to distinguish between competition and no competition. Epitope binning of the five nanobodies showed that they bound in two different epitope bins (Group B and C) (Fig 2E). Group B nanobodies were 3022 and 3023 and Group C nanobodies were 3020, 3024 and 3031 (Fig 2E). Nanobody 3024 showed partial competition with 3020 and 3031, however we still considered this competition and grouped them in the same epitope bin. In addition, all these nanobodies bound to a different site to 2700 (Group A), which demonstrated that our structure prediction-driven selection using an NGS dataset was successful in identifying nanobodies that bind to different sites on PvAMA1 (Fig 2E).

### Nanobodies against PvAMA1 block parasite invasion

We wanted to determine if the PvAMA1 nanobodies could inhibit blood stage invasion. However, since there is no long term *in vitro* culture of *P. vivax*, we used a transgenic W2mef *P. falciparum* parasite line that expresses PvAMA1 (W2-PvAMA1) [38]. Parasite growth was measured via flow cytometry following one replication cycle, and 100% growth was defined as the parasitaemia (percentage of all red blood cells infected) in the absence of nanobody (no Nb, Fig 3A and Fig 3B). We assessed all 14 Sanger-selected PvAMA1 nanobodies (2688 – 2701) at 0.5 mg/mL in growth inhibition assays. These nanobodies were classified either as strong inhibitors (<10% growth, green), intermediate inhibitors (10-80% growth, orange) or non-inhibitors (>80% growth, blue). Five of the nanobodies showed strong inhibition, 2689, 2691, 2699, 2700 and 2701 (Fig 3A). Four nanobodies showed intermediate inhibition, 2688, 2690, 2692 and 2694. The remaining five nanobodies showed no inhibition (Fig 3A). As expected, the negative control SARS-CoV-2 nanobody WNb36 did not inhibit parasite invasion [41].

**Figure 3.**
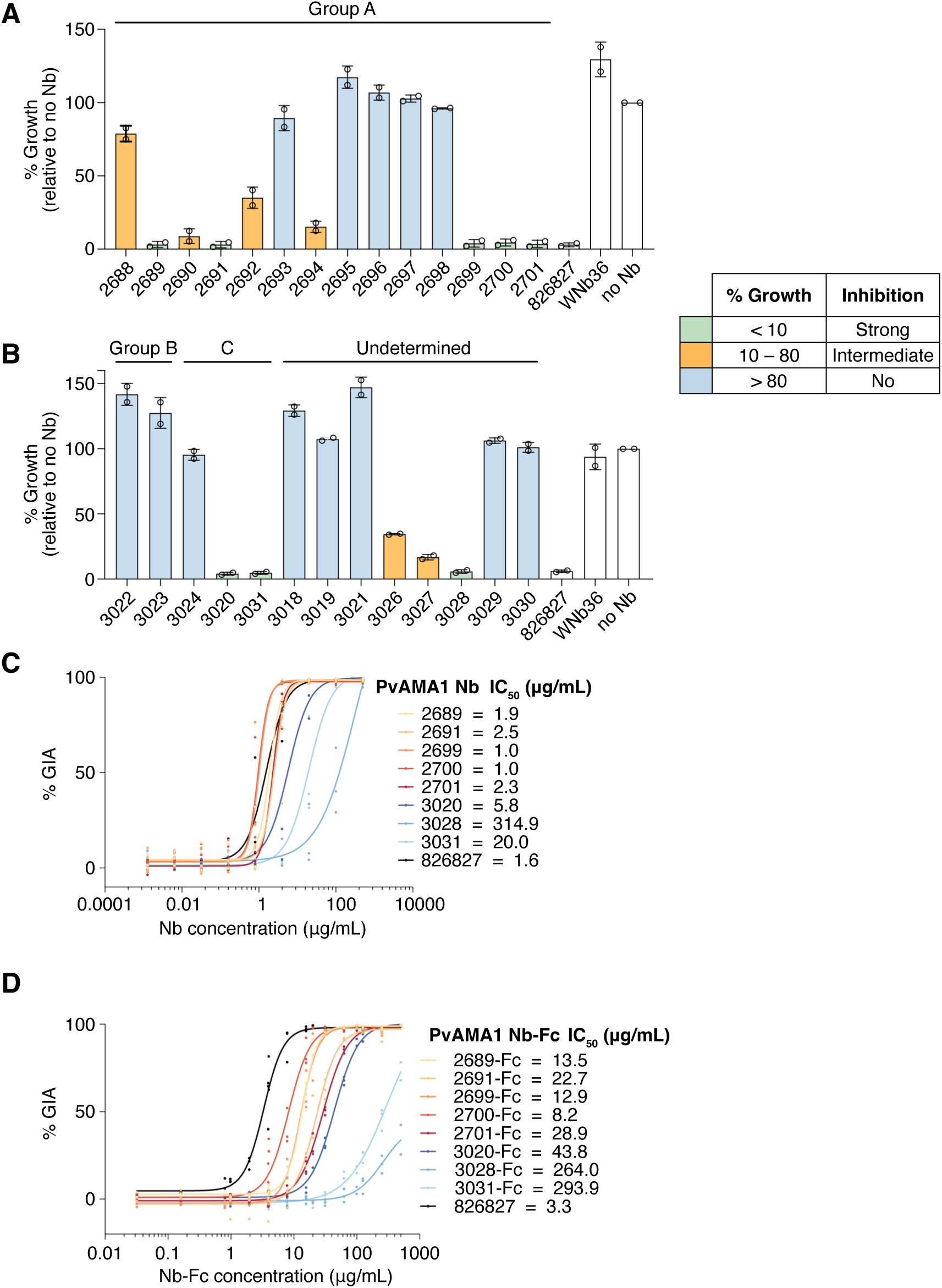
PvAMA1 specific nanobodies and nanobody-Fcs inhibit red blood cell invasion. **(A)** Single concentration growth inhibition assays were performed using Sanger-selected PvAMA1 nanobodies at a final concentration of 0.5 mg/mL using the transgenic *P. falciparum* parasite strain W2-PvAMA1. **(B)** Single concentration growth inhibition assays were performed using NGS and AI-selected nanobodies at a final concentration of 0.5 mg/mL using the transgenic *P. falciparum* parasite strain W2-PvAMA1. Growth inhibitory activity (GIA) IC_50_ values for the inhibitory nanobodies in **(C)** monomeric and **(D)** nanobody-Fc formats, determined through 9-point 1:5 or 1:2 dilutions series ranging from 500 – 0.00128 μg/mL and 250 – 0.98 μg/mL respectively. Resulting data were log transformed and fitted with a 4-parameter non-linear regression curve to calculate the IC_50_ value. Data for A and B are presented as mean ± SD, with each circle representing one independent experiment (n=2 biological replicates). Data for C and D are presented as individual points, representing the result for each concentration per replicate (the number of biological replicates for C, n=3, for D n=2 for 1:5 series and n=3 for 1:2 series). Parasitaemia was measured after one replication cycle. Growth in the absence of nanobody was defined as 100%.

We assessed the 13 AI-assisted selection of NGS nanobodies in growth inhibition assays and categorized them as described above. Two of the Group C nanobodies, 3020 and 3031, and one nanobody in the group with undetermined epitopes, 3028, showed strong inhibition (Fig 3B). Two lower affinity nanobodies with undetermined epitopes, 3026 and 3027, showed intermediate inhibition. The remaining nanobodies showed no inhibitory activity (Fig 3B).

For the strong inhibitors, we observed IC_50_ values ranging from 1.0 – 314.9 μg/mL (Fig 3C and S4 Data). In addition, we fused these nanobodies to a human IgG1 Fc to generate nanobody-Fcs. These nanobody-Fcs bound to PvAMA1 with similar affinities by BLI compared to their monomeric forms, except 2701-Fc that had a *K*_D_ of 0.88 nM compared to its monomeric form with a *K*_D_ of 47.75 nM (S3 Fig, S1 Data and S5 Data). These PvAMA1 nanobody-Fcs showed IC_50_ values ranging from 8.2 – 293.9 μg/mL (Fig 3D and S4 Data). From these results, the most potent nanobodies or nanobody-Fcs were 2700 and 2699. As monomeric nanobodies, 2699 and 2700 both had IC_50_ values of 1.0 μg/mL. In the nanobody-Fc format, 2699-Fc had an IC_50_ of 12.9 μg/mL and 2700-Fc had an IC_50_ of 8.2 μg/mL. The difference in potency between monomeric nanobodies (∼15 kDa) and nanobody-Fc (∼ 80 kDa) are mainly due to the difference in their molecular weights. As a benchmarker, the human monoclonal antibody 826827 showed IC_50_ values ranging from 1.6 to 3.3 μg/mL across the two experiments, which is similar to the previously reported IC_50_ value of 2.6 μg/mL (Fig 3C, 3D) [27].

### Inhibitory nanobodies against PvAMA1 bind conformational epitopes and to a distinct site from 826827

We wanted to determine if the strong nanobody inhibitors bound to the same epitope bin as 826827. We had previously determined Group A inhibitors 2691, 2693 and 2700 bound to a different site from 826827 (Fig 1C). We repeated the epitope binning to include all inhibitory Group A nanobodies (2689, 2691, 2699, 2700 and 2701) and confirmed they bind to a different site from 826827 (Fig 4A). Epitope binning with the Group C inhibitors 3020 and 3031 showed that only 3031 directly competed with 826827 (Fig 4A). We did not include nanobody 3028 due to its low binding signal, which makes it difficult to distinguish between competition and no competition.

**Figure 4.**
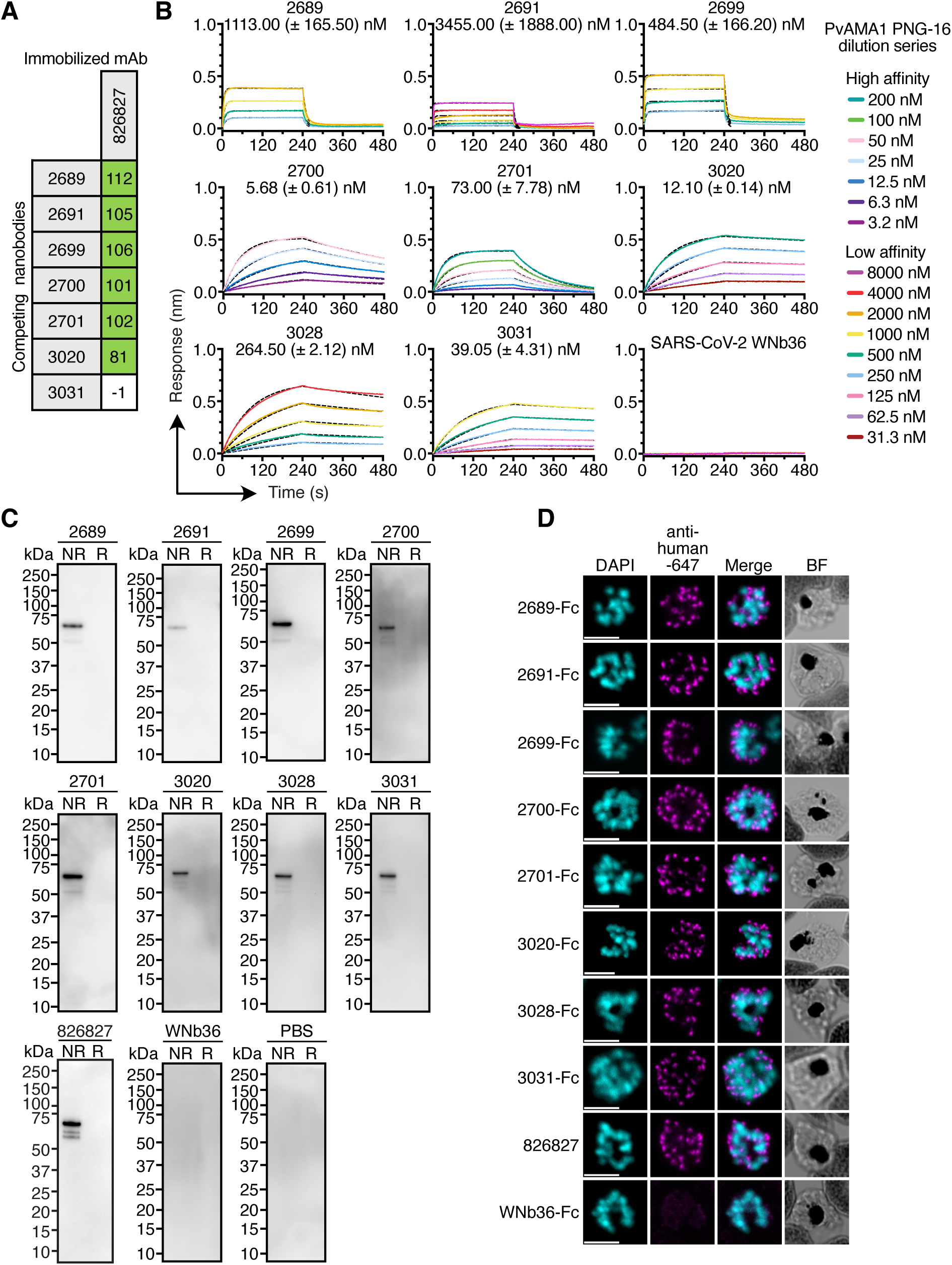
PvAMA1 specific nanobodies recognize PvAMA1 in transgenic W2-PvAMA1 parasites. **(A)** Epitope binning of strong nanobody inhibitors against human monoclonal antibody 826827 using BLI. 826827 was immobilized and dipped into nanobodies pre-incubated with PvAMA1 Palo Alto at a 10:1 molar ratio. Binding of PvAMA1 pre-mixed with nanobody to 826827 was calculated relative to PvAMA1 binding to 826827 alone, which was assigned to 100%. The green and white boxes represent non-competing and competing nanobodies, respectively. **(B)** BLI affinity curves using a two-fold serial dilution of PvAMA1 PNG-16 binding to immobilized nanobodies. Measurements were plotted (solid line) and fitted to a 1:1 binding model (black dashed line). *K*_D_ values represent means with standard deviations from two independent experiments (n=2) and a representative binding curve is shown from one experiment. **(C)** Immunoblots of nanobodies against PvAMA1 from transgenic *P. falciparum* line W2-PvAMA1 schizont lysate. Non-reduced (NR) and reduced (R) lysate was separated by SDS-PAGE and probed with anti-PvAMA1 nanobodies, irrelevant nanobody WNb36 and human monoclonal antibody 826827, and PBS. Anti-His and HRP conjugated anti-mouse IgG were used to detect nanobodies, and HRP conjugated anti-human IgG was used to detect 826827. Molecular weight markers are indicated on the left-hand side of each blot. **(D)** Immunofluorescence assay of schizont stage transgenic *P. falciparum* line W2-PvAMA1 parasites. The indicated nanobody-Fcs and monoclonal antibodies were detected with anti-human conjugated to AlexaFluor 647 (shown in magenta). Parasite nuclei are stained with DAPI (shown in cyan). Merged and brightfield (BF) images are shown. Scale bar = 5 μm.

In addition, we wanted to examine if our strong inhibitory nanobodies would recognize a variant of PvAMA1 called PNG-16, as PvAMA1 is highly polymorphic. The PNG-16 variant is 97% identical to the Palo Alto variant which was used for the alpaca immunisations. However, compared to the Palo Alto variant, PvAMA1 PNG-16 contains 13 of 22 polymorphisms identified to be under balancing selection in *P. vivax* circulating in Papua New Guinea, many of which are located in PvAMA1 D1 in the area surrounding the hydrophobic groove [45]. Using BLI, we observed that the monomeric forms of 2689, 2691 and 2699 showed a 200 to 700-fold reduction in binding affinity to PvAMA1 PNG-16 compared to PvAMA1 Palo Alto, whereas 2700, 2701, 3020, 3028 and 3031 retained affinities within the same magnitude between the two variants (Fig 4B, S1 Fig, S2 Fig, S1 Data and S5 Data). In the nanobody-Fc forms, 2689-Fc, 2691-Fc and 2699-Fc showed >900-fold reduction in binding affinity to PvAMA1 PNG-16 compared to PvAMA1 Palo Alto. 2700-Fc and 2701-Fc bound PvAMA1 PNG-16 with *K*_D_ of 10.51 nM and 83.15 nM, showing 26-fold and 94-fold reduction in binding affinity compared to PvAMA1 Palo Alto, respectively. 3020-Fc, 3028-Fc and 3031-Fc retained similar affinities between 27.90 – 228.50 nM (S3 Fig and S5 Data).

We used immunoblotting to determine whether the inhibitory PvAMA1 nanobodies could detect PvAMA1 in the transgenic W2-PvAMA1 line [38]. In late stage schizonts, PvAMA1 is proteolytically processed from an 81 kDa precursor to the 64 kDa form, which is presented on the merozoite surface for red blood cell invasion [38]. All eight inhibitory nanobodies and 826827 detected a ∼64 kDa band in parasite lysate under non-reducing conditions (Fig 4C) [27]. Under reducing conditions, neither the inhibitory nanobodies nor 826827 showed any reactivity, indicating that their binding to PvAMA1 is dependent on conformational epitopes (Fig 4C). As expected, neither the SARS-CoV-2 nanobody WNb36 nor PBS showed reactivity against PvAMA1 under non-reducing or reducing conditions (Fig 4C) [41].

We conducted an immunofluorescence assay with late-stage schizonts using the inhibitory nanobody-Fcs and 826827, and anti-human IgG conjugated to AlexaFluor 647 for detection. For all the antibodies, we observed punctuate loci localized to the apical end of merozoites, as previously seen for PvAMA1 in late schizont stages (Fig 4D) [38]. The negative control WNb36-Fc did not show any detectable staining of schizonts (Fig 4D). Collectively, these results show that the inhibitory nanobodies could detect PvAMA1 in malaria parasites.

### Inhibitory nanobodies 2700 and 2701 bind PvAMA1 outside the RON2 binding site

To understand the structural basis of nanobody inhibition, we determined the crystal structures of nanobodies 2700 and 2701 bound to PvAMA1 Palo Alto at resolutions of 2.8 and 2.5 Å, respectively (Table 1).

**Table 1:** Data collection and refinement statistics for PvAMA1-864865-2700 and PvAMA1-2701.

|  | **PvAMA1-864865-2700**<br>**(PDB 9ZXJ)** | **PvAMA1-2701**<br>**(PDB 44IE)** |
|---|---|---|
| Data collection statistics |  |  |
| Wavelength (Å) | 0.953725 | 0.953720 |
| Space group | P1 | C222<sub>1</sub> |
| Cell axes (Å) (a, b, c) | 49.9, 74.2, 176.1 | 69.1, 197.0, 180.4 |
| Cell angles (º) (α, γ, β) | 93.3, 91.7, 109.4 | 90, 90, 90 |
| Resolution range (Å) | 47.00-2.78 (2.85-2.78)* | 29.78-2.50 (2.65-2.50)* |
| Completeness (%) | 97.8 (97.8) | 99.8 (99.4) |
| Total no. of reflections | 175,897 (12,604) | 579,758 (94,038) |
| Unique reflections | 58,424 (4,331) | 43,111 (6,822) |
| Redundancy | 3.0 (2.9) | 13.4 (13.8) |
| R<sub>meas</sub> (%) | 16.3 (109.0) | 16.2 (139.5) |
| CC<sub>1/2</sub> (%) | 99.1 (43.2) | 99.9 (81.0) |
| I/σ | 6.76 (1.28) | 13.10 (1.83) |
| Wilson B (Å<sup>2</sup>) | 55.69 | 57.08 |
| Refinement statistics |  |  |
| R<sub>work</sub>/R<sub>free</sub> (%) | 23.0 / 26.9 | 25.6 / 30.6 |
| No. of atoms |  |  |
| Protein | 14,563 | 7,514 |
| Water | 84 | 58 |
| B factors (Å<sup>2</sup>) |  |  |
| Chain A | 56.6 | 62.8 |
| Chain B | 59.5 | 65.3 |
| Chain C | 72.9 | 73.9 |
| Chain D | 74.6 |  |
| Chain E | 53.7 |  |
| Chain F | 76.5 |  |
| Chain G | 55.1 | 44.9 |
| Chain H | 76.1 |  |
| Water | 43.6 | 48.2 |
| R.m.s. deviations |  |  |
| Bond lengths (Å) | 0.003 | 0.009 |
| Bond angles (º) | 0.618 | 1.073 |
| Validation |  |  |
| Ramachandran outliers (%) | 0.00 | 0.00 |
| Ramachandran favored (%) | 96.4 | 96.5 |
| Rotamer outliers (%) | 0.27 | 0.83 |
| C-beta outliers | 0.00 | 0.00 |
| MolProbity score | 1.62 | 1.76 |

\* The values in parentheses represent the highest-resolution shell.

Both nanobodies recognize largely overlapping conformational epitopes on PvAMA1, which is consistent with our epitope binning experiments (Fig 5A-C, S1C Fig). These nanobodies bind PvAMA1 using residues of all three CDR loops, which engage with PvAMA1 at the interface of D1 and D2 (Fig 5C). Nanobody 2700 forms direct interactions with nine PvAMA1 residues, with four residues located in D1 and five residues in D2 (Fig 5D, Table S1). The interaction between 2700 and PvAMA1 results in a buried surface area of 997 Å^2^ (Fig 5D *right panel; outlined in red,* Table S1). Nanobody 2701 interacts with 10 PvAMA1 residues, of which five are located in D1 and D2 respectively, resulting in a buried surface area of 908 Å^2^ (Fig 5D *right panel; outlined in yellow,* Table S2). In both structures, the flexible D2-loop (PvAMA1 residues 296-332) has elevated B-factors (S4 Fig) but is partly visible in its closed position on D1 of PvAMA1, which covers a part of the RON2 binding site (Fig 5A, B and E).

**Figure 5.**
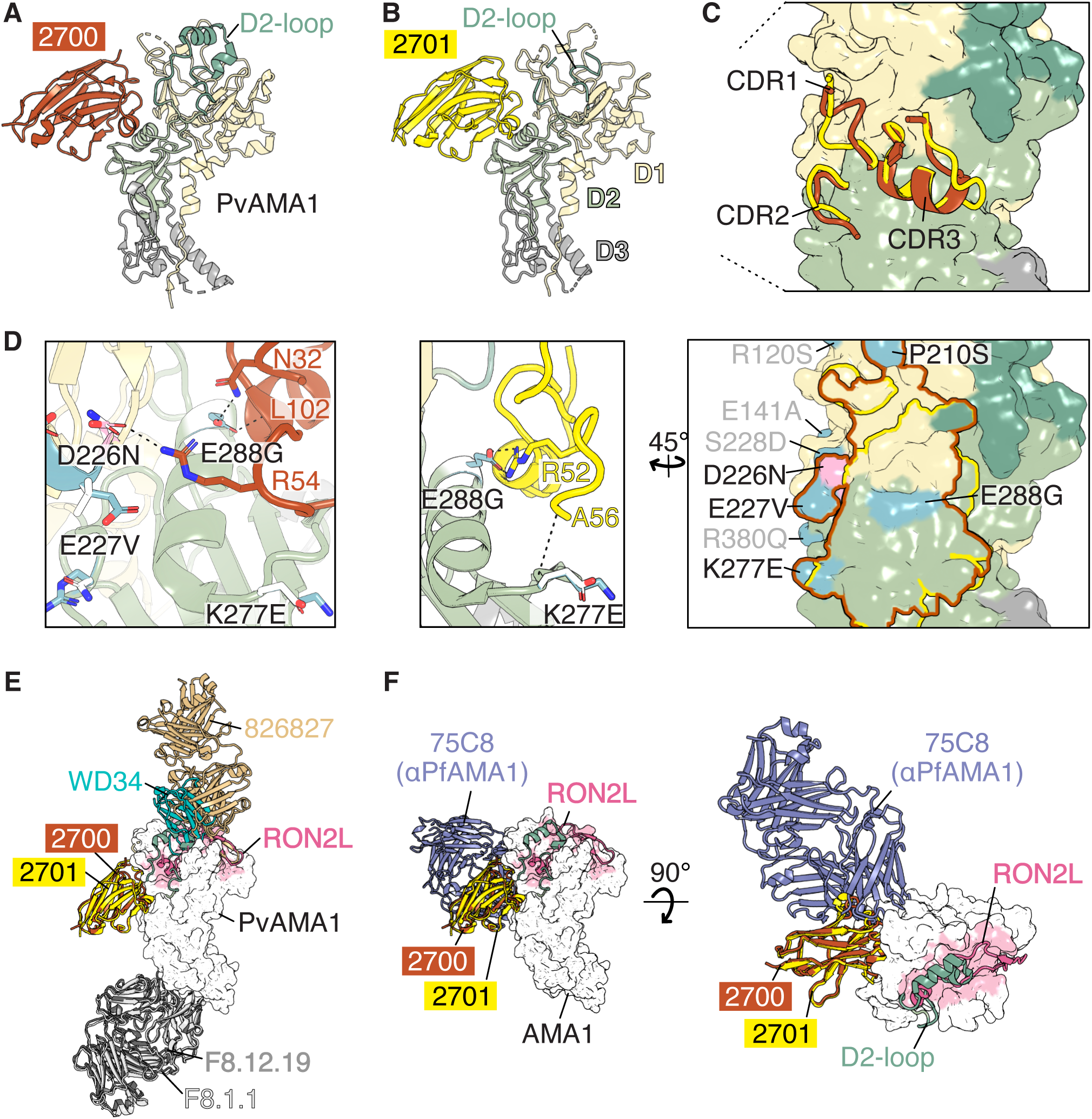
Crystal structures of PvAMA1 in complex with inhibitory nanobodies. Ribbon representations of the PvAMA1-2700 complex (**A**) and PvAMA1-2701 complex (**B**) with 2700 shown in red, 2701 in yellow and PvAMA1 domain 1 in beige (D1, residues 63-248), domain 2 in green (D2, residues 249-385) and domain 3 in grey (D3, residues 386-474). The PvAMA1 D2- loop is indicated. **(C)** All three CDR-loops (CDR1-3) of 2700 and 2701 are involved in binding to PvAMA1. The surface representation of PvAMA1 is coloured as in panels A and B. (**D**) Structural analysis of nanobody interactions with polymorphic residues on PvAMA1. PNG-16 amino acid differences within the epitope are labelled black (P210S, D226N (sequon mutation in Palo Alto construct), E227V, K277E, E288G); Polymorphic residues nearby are labelled in grey. Side chains of polymorphic residues are shown in blue for the Palo Alto variant; a sequon mutation of the Palo Alto expression construct D226N, is shown in pink. Polymorphisms in PNG-16 variant are modelled and indicated in white. CDR loops and residues of 2700 involved in contacting polymorphic residues and the sequon mutation are shown in red (*left*); those of 2701 are shown in yellow (*middle*). Surface of PvAMA1 is coloured as in panel C with epitopes of 2700 and 2701 indicated as outlines, polymorphic residues in blue and sequon mutation in pink (*right*). (**E**) Nanobodies 2700 and 2701 bind distinct sites to the RON2L binding site and to other known Pv-specific antibody epitopes. Overlay of all known crystal structures of PvAMA1 in complex with antibody fragments, with human Fab 826827 (PDB ID 9DX6), i-body WD34 (PDB ID 8QUS), Fab F8.1.1 (PDB ID 8REK), Fab F8.12.19 (PDB ID 2J4W), as well as with RON2L (PDB ID5NQG). All structures were aligned based on PvAMA1. (**F**) Nanobodies 2700 and 2701 bind to a region on PvAMA1 that partly overlaps with where human mAb 75C8 recognizes PfAMA1 (PDB ID 9BJH). PvAMA1-RON2L (PDB ID 5NQG) is depicted as a white surface with RON2L (ribbon representation) and the RON2L binding site coloured in pink. Antibody fragments and PvAMA1 D2-loop of PDB ID 9ZXJ are shown in ribbon representations.

Numerous polymorphisms between PvAMA1 Palo Alto and PNG-16 variants are present on the surface of the protein. The sequence identity between the PvAMA1 expression constructs is 97% with 10 polymorphic residues in D1 and 5 in D2 (S4 Fig). Four polymorphisms, P210S, E227V, K277E and E288G, and one expression construct dependent mutation, D226N (a sequon disruption to avoid glycosylation), are within the epitope recognized by 2700 (Fig 5D *left and right*). Of these, two residues are involved in direct interactions with the nanobody (Fig 5D *left*). D226 of PvAMA1 forms a salt bridge with R54 of 2700. An asparagine at position 226 in the PNG-16 variant could still form a hydrogen bond with the guanidine group of R54 but the ionic electrostatic attraction would be lost, resulting in a weaker interaction. The side chain of E288 forms hydrogen bonds to the side chain of N32 of 2700 and the backbone of L102 (Fig 5D *left*). The change to glycine in the PNG-16 isolate will disrupt these interactions.

For nanobody 2701, two polymorphic PvAMA1 residues, K277 and E288, are involved in direct contacts with the nanobody (Fig 5D *middle panel*). The backbone of K277 forms a hydrogen bond with the backbone of A56 of 2701. The mutation of lysine to glutamic acid in the PNG-16 variant is not expected to affect this backbone-backbone interaction. The side chain of E288 forms a salt bridge with R52 of 2701 and this interaction is lost when E288 is changed to glycine in the PNG-16 variant (Fig 5D *middle panel*). Collectively, the amino acid differences at the nanobody epitopes between the two PvAMA1 variants indicate a minor reduction in the number of interactions and interaction strength for PvAMA1 PNG-16 and may provide an explanation of the slightly lower affinities of 2700 and 2701 to PvAMA1 PNG-16 (Fig 4B, S5 Data) compared to PvAMA1 Palo Alto (Fig 1B, S1 Fig, S1 Data).

The epitopes of nanobodies 2700 and 2701 are distinct to epitopes of previously identified antibodies against PvAMA1 (Fig 5E). The Fab of 826827 and cross-reactive i-body WD34 target epitopes on D1 of PvAMA1 that overlap with the RON2 binding site [27,28]. In contrast, Fabs F8.1.1 and F8.12.19 recognize PvAMA1 D3 [39,40]. We observe no overlap of any of these antibody epitopes with our 2700 and 2701 nanobodies, demonstrating that the nanobodies target novel neutralising epitopes on PvAMA1 (Fig 5E). However, nanobodies 2700 and 2701 do target epitopes on PvAMA1 that have structural overlap with the PfAMA1 region recognized by neutralizing human mAb 75C8 [46] (Fig 5F). While all three inhibitory antibodies recognize epitopes on AMA1 outside the RON2L-binding site, the angle of approach differs between 75C8 Fab and nanobodies 2700 and 2701 (Fig 5F).

### PvAMA1 nanobodies 3020 and 3031 inhibit *P. falciparum* invasion

To assess whether our strong inhibitory PvAMA1 nanobodies can block *P. falciparum* invasion, we used the *P. falciparum* strain 3D7 in a growth inhibition assay. Parasite growth was measured via flow cytometry following one replication cycle, and 100% growth was defined as the parasitaemia in the absence of nanobody (no Nb, Fig 6A). We assessed the eight nanobodies which were shown to be strongly inhibitory against the W2-PvAMA1 line, as well as the NGS nanobodies, as most were predicted to bind close to the conserved hydrophobic groove of AMA1. Nanobodies were assessed at 0.5 mg/mL and classified as either strong inhibitors (<10% growth, green), intermediate inhibitors (10-80% growth, orange) or non-inhibitors (>80% growth, blue). Of these nanobodies, 3020 and 3031 showed strong inhibition and 3027 showed intermediate inhibition against *P. falciparum* (Fig 6A). The remaining nanobodies showed no inhibition (Fig 6A). We had previously shown that 3020 and 3031 are classified as epitope Group C whereas 3027 had an undetermined epitope for PvAMA1 (Fig 2E, Fig 3B). For the strong inhibitors against *P. falciparum* 3D7, we observed IC_50_ values of 2.2 μg/mL for 3020 and 11.9 μg/mL for 3031 nanobodies (Fig 6B and S6 Data). For the nanobody-Fc formats, their IC_50_ values were 16.1 μg/mL for 3020-Fc and 135.2 μg/mL for 3031-Fc (Fig 6B and S6 Data).

**Figure 6.**
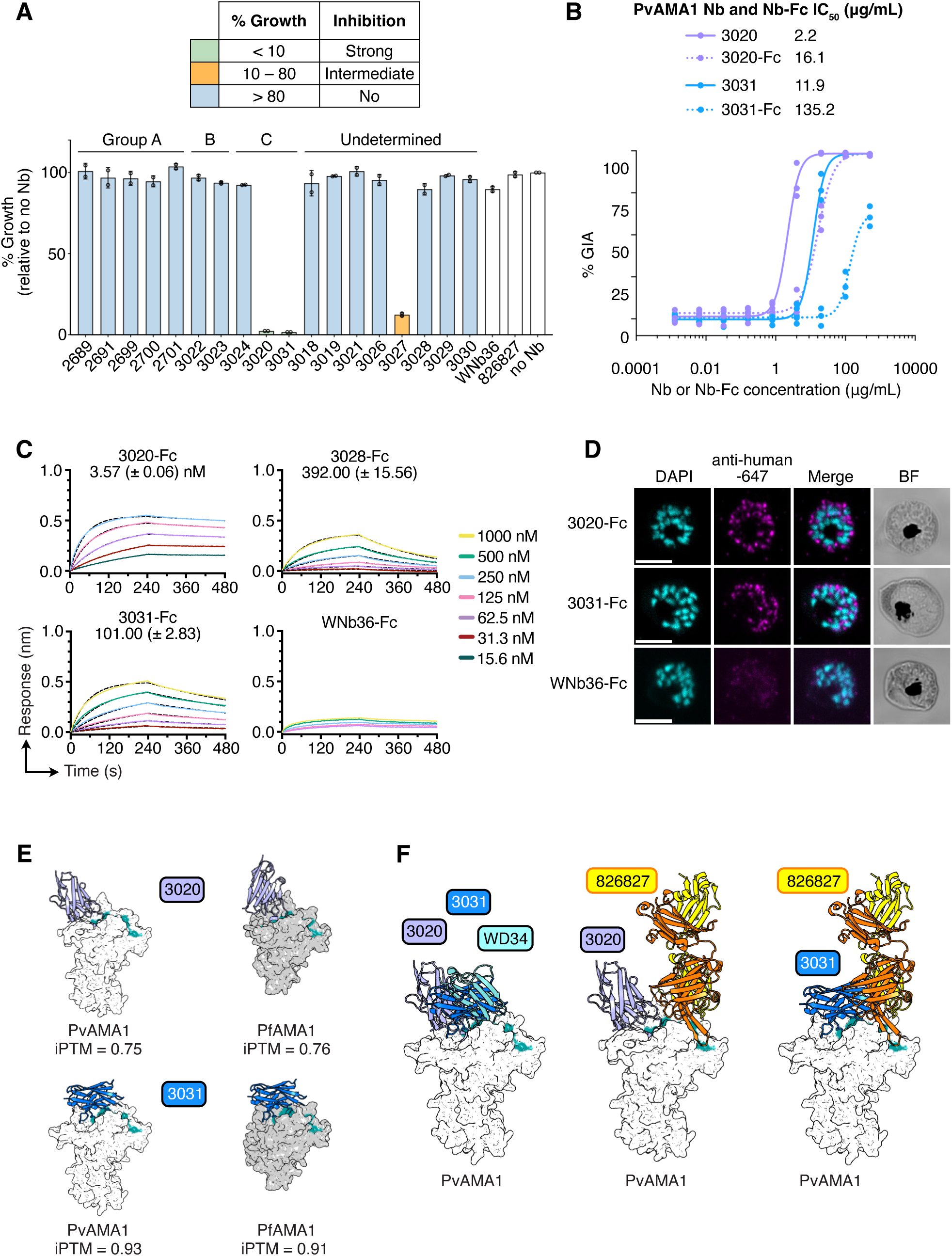
PvAMA1 nanobodies block *P. falciparum* invasion. **(A)** Single concentration growth inhibition assays were performed using the nanobodies at a final concentration of 0.5 mg/mL and the *P. falciparum* strain 3D7. Data is presented as mean ± SD, with each circle representing one independent experiment (n=2 biological replicates). **(B)** Growth inhibitory activity (GIA) IC_50_ values for monomeric nanobodies and nanobody-Fc formats were determined through nine-point 1:5 dilution series ranging from 500 – 0.00128 μg/mL. Resulting data were log transformed and fitted with a 4-parameter non-linear regression curve to calculate the IC_50_ value. Data is presented with individual points representing the result for each concentration per replicate (n=3 biological replicates). Parasitaemia for A and B was measured after one replication cycle. Growth in the absence of nanobody was defined as 100%. **(C)** BLI affinity curves using a two-fold serial dilution of PfAMA1 binding to immobilized nanobody-Fcs. Measurements were plotted (solid line) and fitted to a 1:1 binding model (black dashed line). *K*_D_ values represent means with standard deviations from two independent experiments (n=2) and a representative binding curve is shown from one experiment. **(D)** Immunofluorescence assay of schizont stage *P. falciparum* strain 3D7 parasites. The indicated nanobody-Fcs were detected with anti-human conjugated to AlexaFluor 647 (shown in magenta). Parasite nuclei are stained with DAPI (shown in cyan). Merged and brightfield (BF) images are shown. Scale bar = 5 μm. **(E)** Structural predictions of 3020 and 3031 binding to PvAMA1 and PfAMA1. 3020 is shown in purple (*top*) and 3031 in blue (*bottom*), PvAMA1 is shown as a white surface and PfAMA1 as a grey surface with the hydrophobic grooves of domain 1 colored in cyan. Protenix iPTM scores are indicated. **(F)** Structural overlays of the PvAMA1-predictions with 3020, 3031 and the crystal structures of ibody WD34 (left), and 826827 (*middle*, and *right*). 3020, 3031 and PvAMA1 are depicted as in panel F. WD34 is shown in light blue and 826827 in orange and yellow (heavy and light chains respectively).

Using BLI, we determined that of the eight nanobody-Fcs which bind PvAMA1, only 3020-Fc, 3028-Fc and 3031-Fc showed binding to PfAMA1 (S5 Fig). Affinities were between *K*_D_ 3.57 and 392 nM for the three nanobody-Fcs that bound to PfAMA1 (Fig 6C and S7 Data). We conducted an immunofluorescence assay with late-stage schizonts using the strong inhibitory nanobody-Fcs and anti-human IgG conjugated to AlexaFluor 647 for detection. For both 3020-Fc and 3031-Fc we observed punctate loci localized to the apical end of merozoites (Fig 6D). The negative control WNb36-Fc did not show any detectable staining of schizonts (Fig 6D) [41]. Collectively, these results show that the PvAMA1 nanobodies 3020 and 3031 recognize PfAMA1 and can potently inhibit *P. falciparum* invasion.

We were unsuccessful in obtaining crystal structures of 3020 and 3031 with PvAMA1. Our initial structural predictions for NGS nanobodies binding to PvAMA1 were done using AlphaFold 3, however, we used Protenix to redo the predictions for nanobodies 3020 and 3031 for binding to PfAMA1. The Protenix model has high-confidence for nanobody 3031 binding to PvAMA1 and PfAMA1 with iPTM scores > 0.9 (Fig 6E). Structural predictions of 3020 binding to PvAMA1 and PfAMA1 were of lower confidence with iPTM scores of 0.75 and 0.76 respectively (Fig 6E). The structural predictions suggest that 3020 and 3031 nanobodies bind to the hydrophobic groove of D1 of AMA1, the part that is usually covered by the flexible D2 loop. Most predicted interactions are formed with relatively conserved residues of both *Plasmodium* species (S5 Fig). When compared to the crystal structure of i-body WD34, both 3020 and 3031 would have partly overlapping epitopes with WD34 for binding to PvAMA1 (Fig 6F, *left*). The structural predictions are also in agreement with our epitope binning experiments in several ways. First, nanobody 3031 binds to an overlapping epitope with human antibody 826827 (Fig 4A, Fig 6F, *right*), but 3020 does not compete with 826827 (Fig 4A, Fig 6F, *middle*). Second, both nanobody 3020 and 3031 have overlapping epitopes (Fig 2E, Fig 6F, *left*).

## Discussion

Since the discovery that the interaction between AMA1 and RON2 is essential for tight junction formation during malaria parasite invasion of red blood cells, studies have focused on antibodies that block AMA1-RON2 binding. Here, we identified inhibitory nanobodies that target novel epitopes on PvAMA1 using phage display combined with Sanger sequencing, NGS and structure prediction approaches. We show that PvAMA1 nanobodies can potently inhibit parasite invasion *in vitro,* are capable of cross species recognition, and we provide structural insights to their binding epitopes.

In our growth assays, we observed that nanobody 2700 in monomeric format or as nanobody-Fc showed the highest potency in blocking invasion of the W2-PvAMA1 line. Nanobody 2700 had an IC_50_ of 1.0 µg/mL in monomeric nanobody format and 8.2 µg/mL as a nanobody-Fc fusion. Notably, dimerization of the nanobody via the Fc-domain did not enhance neutralization, as the molar IC_50_ values are comparable. In comparison, IC_50_ values of human monoclonal antibody 826827 were 1.6 and 3.3 µg/mL in these assays, demonstrating that the potency of nanobody 2700 is in a similar range. In our collection of nanobodies, nanobody 2700 has the highest binding affinity for PvAMA1, with a *K*_D_ value of 1.7 nM for the Palo Alto variant against which it was raised, and 5.7 nM for the PNG-16 variant. As a nanobody-Fc fusion, 2700 has a *K*_D_ of 0.4 nM for PvAMA1 Palo Alto compared to a *K*_D_ of 10.5 nM for PvAMA1 PNG-16, which represents a 26-fold reduction in affinity but is still considered high affinity binding.

Our epitope binning experiments show that we have inhibitory nanobodies that bind to different sites on PvAMA1. The generation of bi-and tri-paratopic nanobodies can greatly enhance the efficacy of distinct nanobodies as shown for nanobodies against Influenza [47] and SARS-CoV-2 [48–52]. Fusion of nanobodies into multivalent reagents enhanced neutralisation capabilities significantly compared to their monomeric counterparts. Targeting distinct sites simultaneously reduces the risk of escape mutants emerging and resistance developing, and can broaden antigenic breadth [48,49,51,52]. In future studies, combining 2700 with inhibitory nanobodies that bind to nonoverlapping epitopes on PvAMA1, namely Group C nanobodies 3020 and 3031, could further enhance their potencies in growth inhibition, reduce resistance and provide strain-transcending capabilities.

RON2 binds AMA1 within a hydrophobic groove in D1. Antibodies that inhibit *P. falciparum* parasite growth which were shown to bind in or adjacent to the hydrophobic groove include: 1F9, IgNAR, humAbAMA1, and the cross-reactive i-body WD34 [25,26,28,37]. Several antibodies targeting the PvAMA1 hydrophobic groove have been characterized, which include WD34 and the human monoclonal antibody 826827 [27,28]. Inhibitory antibodies which bind AMA1 outside of the hydrophobic groove have also been identified: 4G2, which binds the flexible PfAMA1 D2-loop that is displaced upon RON2 binding [53,54], 75C8, which binds at the base of loops formed by PfAMA1 D1 and D2, and 75B10, which binds PfAMA1 D1 and D2 at the base of the RON2 binding site [46].

We determined the structure of two growth inhibitory nanobodies, 2700 and 2701, in complex with PvAMA1 and show they bind to a novel neutralising epitope on PvAMA1 distinct from the RON2 binding site and epitopes of known antibodies against PvAMA1; 826827, WD34, F8.12.19 and F8.1.1 [27,28,39,40]. The nanobody epitopes overlap with that of inhibitory

PfAMA1 antibody 75C8 [46], however, it should be noted that 75C8 does not bind to PvAMA1. As expected, nanobodies 2700 and 2701 did not bind PfAMA1, as their epitopes in PvAMA1 differ in sequence from PfAMA1. These epitopes are outside of the RON2 binding site and the antibody/nanobodies are unlikely to sterically interfere with AMA1-RON2 interaction. Antibody 75C8 has been proposed to prevent the displacement of the D2-loop, however the D2-loop is not visible in the crystal structure of the PfAMA1-75C8 complex (PDB ID 9BJH) and so further investigation may be needed to understand its mode of inhibition [46]. In our crystal structures, the D2-loop is partly visible in the cleft-bound position on D1 of PvAMA1 but is only involved in one contact with 2700 (Y298 of the PvAMA1 D2-loop forms a hydrogen bond with E111 of 2700). Based on the location of the nanobody epitopes and only one interaction with a residue at the base of the D2-loop, we do not expect the nanobodies to prevent the PvAMA1 D2-loop from moving into the open conformation required for RON2 binding. Prior studies have suggested that other regions of AMA1, outside of the hydrophobic groove, may be involved in binding interactions for invasion [55], however this has not been confirmed.

Antigen polymorphisms can limit neutralization capabilities of antibodies and nanobodies and PvAMA1 is a highly polymorphic surface antigen [45,56,57]. To determine the specificity of our nanobodies, we tested their binding to recombinant PvAMA1 proteins from two different *P. vivax* strains; Palo Alto, the variant used for alpaca immunization and PNG-16, which has significant sequence polymorphisms compared to Palo Alto [27,45]. While some nanobodies did show a reduced affinity to PvAMA1 PNG-16 compared to PvAMA1 Palo Alto, we confirmed that alongside nanobody 2700, the inhibitory nanobodies 2701, 3020, 3028 and 3031 retained similar, albeit reduced, affinities for the PNG-16 variant in agreement with our structural analysis.

Our initial nanobody panel, 2688 – 2701, was identified using Sanger sequencing and all bound within the same epitope bin (Group A), which was distinct from the binding site of 826827 which overlaps with the hydrophobic groove. To select nanobodies that bind to different sites on PvAMA1 from Group A nanobodies, we analysed our NGS nanobody dataset using AlphaFold 3 and performed structure predictions to identify those likely to bind PvAMA1 within the hydrophobic groove where RON2 binds. Characterisation of these NGS selected nanobodies showed some consistency with the structural predictions. First, five nanobodies identified using NGS and structure prediction approaches, 3020, 3022, 3023, 3024 and 3031, bound to two different epitope bins distinct from the Group A nanobodies. Second, nanobodies 3022 and 3023 compete with each other, showing that they bind to overlapping epitopes, which is consistent with their structural predictions. Third, nanobody 3031 competes with 826827, which is known to partially occupy the hydrophobic groove of PvAMA1 [27].

However, we observed some discrepancies between the AlphaFold 3 structural predictions and the experimental data for the NGS selected nanobodies against PvAMA1. First, 3020 should compete with 826827, but we did not observe any competition between these two antibodies. Second, structure predictions suggest that 3024 has a distinct binding site to nanobodies 3020 and 3031. However, we did observe some binding competition between 3024 with these nanobodies. Despite the discrepancies, these results show that the use of the NGS dataset combined with structure prediction approaches was able to identify nanobodies that bind outside of the Group A epitope, and identified nanobody 3031 which is likely to bind to the hydrophobic groove of PvAMA1.

It has previously been shown that the transgenic line W2-PvAMA1 can be inhibited by targeting the hydrophobic groove with the i-body WD34 [28]. However, the mechanism of inhibition is unresolved. Data suggests that PvAMA1 expressed in the *P. falciparum* transgenic line has reduced binding of PfRON2, and these parasites are not inhibited by PfRON2 peptides [55,58]. In this study, we showed that nanobody 3031 binds a similar epitope of PvAMA1 to 826827, which partially occupies the hydrophobic groove, and that this nanobody inhibits parasite invasion with an IC_50_ value of 20.0 μg/mL. These results support a functional role for the hydrophobic groove of PvAMA1 in this transgenic line in red blood cell invasion, which may be independent of RON2 binding.

We also showed that nanobodies 3020 and 3031 were strong inhibitors of *P. falciparum* invasion, with IC50 values of 2.2 and 16.1 µg/mL, respectively. To the best of our knowledge, the i-body WD34 is the only antibody-like molecule with similar *in vitro* inhibitory activity against both *P. vivax* and *P. falciparum* AMA1 [28], as the PvAMA1 specific antibody F8.12.19 showed significant cross-species reactivity but had no major inhibitory activity *in vitro* [39]. In addition, 3020-Fc binds with a higher affinity to PfAMA1 (*K*_D_ of 3.57 nM) when compared to PvAMA1 (*K*_D_ of 21.75). Although we were unable to obtain crystal structures of either 3020 or 3031 with *Plasmodium* AMA1, structure predictions using Protenix suggest that they may recognise relatively conserved residues within the central region of the hydrophobic groove in both PvAMA1 and PfAMA1. However, further structural studies are required to confirm their binding sites.

Nanobodies have been identified to block invasion and transmission of *Plasmodium* species but most of these studies have focused on *P. falciparum* [59–64]. To the best of our knowledge, we have characterized the first collection of nanobodies against PvAMA1 that block invasion. Our findings demonstrate that we can block both PvAMA1 and PfAMA1 function in parasite invasion using nanobodies, and that PvAMA1 has invasion-inhibitory epitopes within and outside of the hydrophobic groove. This knowledge could inform the design of vaccines based on

PvAMA1 that induce antibodies with potent inhibitory activity and circumvent the potential for vaccine escape due to sequence polymorphisms. Additional structural studies are required to understand how other inhibitory PvAMA1 nanobodies block parasite invasion to fully understand the functional landscape of PvAMA1. Future experiments using *P. vivax* clinical isolates are required to further define the efficacy of these nanobodies for inhibition of invasion of *P. vivax* into reticulocytes.

## Materials and Methods

### Expression and purification of PvAMA1

PvAMA1 was expressed using a previously described expression construct (ACB42433.1; Palo Alto genotype) [27,55]. The construct consists of a signal peptide for tissue plasminogen activator followed by a 6xHis-tag and sequence coding for PvAMA1 residues G22-Q484. PvAMA1 was expressed in Expi293 cells via transient transfection at a density of 2.5 × 10^6^ cells/mL following the manufacturer’s protocol. Five days post transfection, the secreted protein was harvested by centrifugation at 7000 x g and dialyzed overnight into 30 mM Tris-HCl pH 7.5, 300 mM NaCl. The supernatant was applied to a 5 mL HisExcel HP column (Cytiva) and the flow through was collected. While His-tagged PvAMA1 was eluted using 100 mM imidazole, 30 mM Tris-HCl pH 7.5, 300 mM NaCl, we observed that the majority of PvAMA1 protein was present in the flow through fraction which migrated at a reduced size compared to His-tagged PvAMA1. The identity of untagged PvAMA1 in the flow through was confirmed using mass spectrometry analysis. The flow through fraction was concentrated and the final purification step for untagged PvAMA1 was performed using a Superdex 200 increase size exclusion chromatography column, which was pre-equilibrated in 20 mM HEPES pH 7.5, 150 mM NaCl.

To reduce flexibility and for crystallization purposes, PvAMA1 coding for residues G42-Y474 was subcloned into a modified version of pTwist-CMV via AgeI and KpnI restriction enzyme sites. The construct consists of the signal peptide of chicken receptor protein-tyrosine phosphatase sigma (RPTPσ) for secretion followed by the PvAMA1 sequence and a non-cleavable C-terminal StrepII-tag. The protein was expressed in Expi293 cells via transient transfection at a density of 2.5 × 10^6^ cells/mL following the manufacturer’s protocol and harvested five days post transfection by centrifugation at 7000 x g. The supernatant was dialyzed for 4 hours at 4°C into 30 mM Tris-HCl pH 7.5, 300 mM NaCl. After sterile filtration, 2 mL of StrepTactin XT resin (IBA) was added and incubated for 30 min on rollers at room temperature. The mix was added to a gravity flow column, washed with 50 mL of 30 mM Tris-HCl pH 7.5, 300 mM NaCl buffer and eluted with 50 mM biotin, 30 mM Tris-HCl pH 7.5, 300 mM NaCl. StrepII-tagged PvAMA1 was concentrated and further purified by Superdex 200 increase 10/300 size exclusion chromatography column (Cytiva) pre-equilibrated in 20 mM HEPES pH 7.5, 150 mM NaCl.

### Isolation of nanobodies against PvAMA1

One alpaca was immunized six times with ∼300 µg of recombinant PvAMA1 using GERBU FAMA as adjuvant. Immunization and handling of the alpaca for scientific purposes was approved by Agriculture Victoria, Wildlife & Small Institutions Animal Ethics Committee, project approval No. 36.20. Blood was collected three days after the last immunization for the preparation of lymphocytes. Nanobody library construction was carried out according to established methods [64]. Briefly, alpaca lymphocyte mRNA was extracted and amplified by RT-PCR with specific primers to generate a cDNA library. The library was cloned into a pMES4 phagemid vector, amplified in TG1 *E. coli* cells and subsequently infected with M13K07 helper phage (NEB, Cat # N0315) for phage expression. Phage display was performed as previously described [62,64] with two rounds of biopanning on 1 µg of immobilized PvAMA1. Positive hits were identified using ELISA and sequenced using Sanger sequencing and annotated using IMGT/V-QUEST [65] and aligned in Geneious Prime 2022.1.1.

### Next-generation sequencing sample preparation of nanobody phage libraries

Plasmids from phage libraries were digested and gel purified to isolate the nanobody sequences. Paired-end 2x300 bp sequencing libraries were prepared using the NEBNext Multiplex Oligos for Illumina (Cat # E7395) in a PCR-free manner according to the manufacturer’s instructions. The samples were size selected using AMPure XP beads, quantified by qPCR using the KAPA Library Quantification Kit and sequenced on an Illumina NextSeq 2000 instrument. Raw data processing and most analysis was performed using Alpseq v0.5.0 [42] with default settings. Cladograms were generated from generalized Levenshtein distances between CDR3s with the Neighbor-Joining method [66] from the phangorn (v2.12.1) [67] package and plotted using the ggtree (v3.14.0) [68] package in R (v4.4.1) [69].

### AI-assisted selection of PvAMA1 nanobodies

Structural prediction of nanobody–PvAMA1 complexes was performed using AlphaFold 3 [43] on the Walter and Eliza Hall Institute of Medical Research (WEHI) high-performance computing cluster. Predictions were performed using four independent model seeds, with five diffusion samples generated per seed. The highest-confidence docking poses for each complex were manually inspected in PyMOL [70] to classify predicted binding sites on PvAMA1 and assess the structural plausibility of their interactions.

To estimate complex interaction energies, all twenty models for each complex were analysed using the empirical force field FoldX [44]. Prior to energy calculation, each predicted complex was subjected to five successive rounds of the FoldX *RepairPDB* command to optimize side-chain conformations. Interaction energies were then calculated using the FoldX *AnalyseComplex* command.

Structural predictions of *P. falciparum* cross-specific nanobodies 3020 and 3031 were performed using Protenix [71] on the WEHI high-performance computing cluster. Predictions with PfAMA1 and for comparison with PvAMA1 were performed using 20 independent model seeds, with five diffusion samples generated per seed and the highest confidence docking poses used for Figure 6. Overall, predictions of 3020 with both PfAMA1 and PvAMA1 were of lower confidence (max iPTM scores = 0.75 and 0.76 and mean iPTM scores = 0.47 and 0.51 for PvAMA1 and PfAMA respectively) than 3031 with the respective AMA1 protein structures (max iPTM scores = 0.93 and 0.91; mean iPTM scores = 0.92 and 0.39 for PvAMA1 and PfAMA respectively).

### Nanobody expression and purification

Nanobodies were expressed in WK6 *E. coli* cells. Bacteria were grown in Terrific Broth supplemented with 0.1% (w/v) glucose and 100 µg/mL carbenicillin at 37°C to an OD_600_ of 0.7, induced with 1 mM IPTG and grown overnight, 180 rpm at 28°C. Cell pellets were harvested and resuspended in 20% (w/v) sucrose, 10 mM imidazole pH 7.5, 150 mM NaCl PBS and incubated on ice for 15 min. EDTA pH 8.0 was added to a final concentration of 5 mM and incubated on ice for 20 min. After this incubation, 10 mM MgCl_2_ was added, and periplasmic extracts were harvested by centrifugation at 4,000 x g for 1 h. The supernatant was sterile filtered with a 0.22 µm filter loaded onto a 1 mL HisTrap Excel column (Cytiva) in 5 mM imidazole pH 7.5, 100 mM NaCl PBS. Nanobodies were eluted with 400 mM imidazole pH 7.5, 100 mM NaCl PBS, and subsequently concentrated and buffer exchanged into 20 mM HEPES pH 7.5, 150 mM NaCl.

### Biolayer Interferometry for affinities

Nanobody affinity measurements were performed using the Octet RED96e instrument (FortéBio). Assays were performed at 25°C in solid black 96-well F-bottom plates (Greiner-Bio One #655209) agitated at 1,000 rpm. Kinetic buffer consisted of PBS with 0.1% (w/v) BSA and 0.05% (v/v) Tween-20. A 60 s baseline step was applied before nanobodies were loaded onto His1K biosensors (Sartorius) by submerging sensor tips in 5 µg/mL of each nanobody until a response of 0.5 nm was obtained. Sensors were then dipped in kinetic buffer for 60 s. Association measurements were performed for 240 s using a two-fold serial dilution of PvAMA1 Palo Alto untagged and dissociation was measured in kinetic buffer for 240 s. For higher affinity nanobodies, the starting concentration of the dilution series were between 100 – 400 nM whereas for lower affinity nanobodies, starting concentrations were between 1000 – 8000 nM. Sensors were regenerated using a cycle of 5 s in 100 mM glycine pH 1.5 and 5 s in kinetic buffer repeated five times. Baseline drift was corrected by subtracting the response of a nanobody-loaded sensor dipped in kinetic buffer for the association step. Curve fitting analysis was performed with Octet Data Analysis 11.1 software using a global fit 1:1 model to determine *K*_D_ values and kinetic parameters. For lower affinity nanobodies with very fast dissociation rates, only the first 20 seconds of data in the dissociation step were used in the analysis. Curves that could not be fitted well were excluded from the analyses. We determined a good fit as R^2^ >0.98 and X^2^ <0.5.

PvAMA1 PNG-16 nanobody affinities were conducted using the same parameters as the above method. Nanobody-Fc affinities to PvAMA1 Palo Alto and PNG-16 were measured with AHC biosensors (Sartorius) using 180 s association and 180 s dissociation, except for 3020-Fc that was measured using 180 s association and 300 s dissociation.

Single binding experiments were performed with nanobody-Fcs immobilized to AHC biosensors and dipped into a single concentration of PfAMA1 at 2000 nM. Nanobody-Fc affinities to PfAMA1 were then measured using a two-fold serial dilution starting at 500 nM or 1000 nM and with 240 s association and 240 s dissocation.

### Biolayer Interferometry for epitope binning

Nanobody epitope binning was performed using a premix setup where 500 nM PvAMA1 Palo Alto was pre-incubated with each nanobody at a 6-fold molar excess (3000 nM) for 1 hour at room temperature. A 30 s baseline step was established between each step of the assay. His1K sensors were first loaded with 10 µg/mL of nanobody for 5 min. The sensor surface was quenched by dipping into 10 µg/mL of an irrelevant nanobody (WNb36 against SARS-CoV-2) for 5 min. Nanobody-loaded sensors were then dipped into premixed solutions of PvAMA1 Palo Alto and nanobody for 5 min. Nanobody loaded sensors were also dipped into PvAMA1 alone to determine the level of PvAMA1 binding to immobilized nanobody in the absence of other nanobodies. Percentage competition was calculated by dividing the maximum response of the premixed PvAMA1 and nanobody solution binding by the maximum response of PvAMA1 binding alone, multiplied by 100. Competition was defined as < 40% binding.

Epitope binning of nanobodies against monoclonal antibody 826827 was performed using a sandwich setup or premix setup. In the sandwich setup, nanobodies were loaded onto a NTA biosensor (Sartorius) at 5 µg/mL for 5 min. After a baseline in buffer, the nanobody loaded sensors were then dipped into PvAMA1 Palo Alto at 100 nM for 5 min. After another baseline, sensors were then dipped into 826827 at 100 nM for 5 min. A positive binding response of 826827 indicates it binds a non-overlapping epitope to the nanobodies, and no binding of 826827 indicates it binds an overlapping epitope with the nanobodies. In the premix setup, 826827 was loaded onto AHC sensors and dipped into a premix of PvAMA1 Palo Alto at 50 nM and nanobody at 10-fold molar excess (500 nM). Percentage competition was calculated as described above.

### Growth inhibition assays

Growth inhibition assays were performed using the previously characterized *P. falciparum* parasite line which expresses PvAMA1 in the W2mef background (called W2-PvAMA1) [38] or the *P. falciparum* strain 3D7. Parasite culture was maintained using O^+^ red blood cells (Australian Red Cross, Lifeblood) at 4 % haematocrit in RPMI 1640, GlutaMAX, HEPES (Thermo Fisher) supplemented with: hypoxanthine (12.5 mg/L) (Sigma), D-(+)-glucose (2 g/L) (Thermo Fisher), gentamycin (20 mg/L) (Thermo Fisher), 0.5 % (w/v) AlbuMAX II (Thermo Fisher). Assays were set up using trophozoite stage parasites at 1 % parasitemia, 2 % haematocrit in 96-well U bottom plates. Assay volumes were 50 μL per well, including 5 μL of relevant nanobody, nanobody-Fc or control treatment. Nanobodies were assessed at 0.5 mg/mL for single concentration assays, 500 – 0.00128 µg/mL in the 1:5 dilution series, or 250 – 0.98 µg/mL in the 1:2 dilution series for IC_50_ titration assays. Plates were incubated at 37°C, in 1% O_2,_ 5% CO_2_, balance N_2_ for ∼48 hours to allow a single replication cycle to proceed. All wells were fixed using 0.125% glutaraldehyde, then stained with 1X SYBR Green I (Invitrogen) diluted in PBS, and parasitaemia was determined using either the Novocyte Quanteon Flow Cytometer (Agilent), or Cytoflex Flow Cytometer (BD). For each well, a total of 9 x 10^4^ red blood events were collected (gated on SSC-A vs. FSC-A, then again on FSC-H vs FSC-A). Parasitized cells were gated using the B530 detector (FITC) vs. FSC-A. Parasite growth was calculated with the following equation: Growth (% PBS/no nanobody) = (parasitemia % treated/parasitemia % control)*100. Growth inhibitory activity was calculated with the following equation: % GIA = 100 – ([parasitemia % treated/parasitemia % control] *100). Samples were tested in technical duplicates, and the number of biological replicates completed is indicated in the figure legend for each experiment. Use of human blood for parasite culturing was approved by the Walter and Eliza Hall Institute Human Research Ethics Committee, project approval No. 86-17.

### Immunoblotting

For immunoblotting, synchronized late schizont stage cultures of W2-PvAMA1 or *P. falciparum* strain 3D7 were used. Infected red blood cell pellets were treated with 0.075% saponin and diluted 1:40 in 2X non-reducing buffer, 10 µL of which was loaded onto a 4-12% Bis-Tris gel under non-reducing and reducing conditions. Blots were probed with 5 µg/mL nanobodies or 0.1 µg/mL of human monoclonal antibody 826827 or PBST for 1 h at RT. Nanobodies were detected using mouse anti-His (Bio-Rad) [1:1000] followed by an anti-mouse IgG-HRP (Merck) [1:1000]. 826827 was detected using anti-human IgG-HRP [1:1000] (Jackson ImmunoResearch). All nanobodies and human antibody were diluted in 1% (w/v) skim milk made up in PBST. Chemiluminescence was detected using SuperSignal West Pico PLUS Chemiluminescent Substrate (Thermo Scientific) and imaged using a ChemiDoc imaging system (Bio-Rad).

### Immunofluorescence assay

Thin blood smears of W2-PvAMA1 or *P. falciparum* strain 3D7 schizont stage parasites were fixed in equal parts of ice-cold acetone and methanol for 10 min and allowed to air dry. The slides were incubated with 3% (w/v) bovine serum albumin (BSA)/PBS for 1 h then washed three times in 1x PBS prior to addition of the 826827 antibody or nanobody-Fcs, which were diluted in 3% BSA/PBS. Primary antibodies were incubated on the slide for 1 h within a humidified chamber. After washing as described, anti-human IgG conjugated to Alexa Fluor 647 (Jackson ImmunoResearch) (diluted 1:1000 in 3% BSA/PBS) was used as a secondary antibody and incubated for 1 h within a humidified chamber. The slides were mounted with a coverslip using VECTASHIELD PLUS antifade mounting medium with DAPI (Vector Laboratories) and sealed with nail polish. Images were acquired on a Zeiss LSM 980 using confocal mode with a 63x (1.4NA) objective lens with oil immersion using the 405, and 639 lasers. Images were processed using FIJI ImageJ software (v2.14.0/1.54) [72] and the Z-projection max intensity images were used in the figures.

### Crystallization and structure determination

Stable complexes were formed between StrepII-tagged PvAMA1 and 2701, and between untagged PvAMA1, 2700 and the Fab fragment of human monoclonal antibody 864865. The complexes were purified by size exclusion chromatography in 20 mM HEPES pH 7.5, 150 mM NaCl before crystallization. Crystallization screens were set up with 4 and 8 mg/mL at 20°C at the Monash Macromolecular Crystallisation Platform (MMCP), Melbourne. Crystals of PvAMA1-2700-864865 were obtained in 10% (w/v) polyethylene glycol (PEG) 6000, 0.1 M tri-sodium citrate pH 5.5, 2% (v/v) glycerol using hanging drop vapour diffusion. One crystal was harvested with 25% (w/v) glycerol in mother liquor and used for data collection. Initial PvAMA1- 2701 crystals grew in 12% (w/v) PEG3350, 4% (v/v) Tacsimate and were optimised in a hanging drop vapour diffusion experiment. A PvAMA1-2701 crystal grown in 8% (w/v) PEG3350, 4% (v/v) Tacsimate was harvested with 30% (v/v) ethylene glycol in mother liquor as cryo-protectant and used for data collection.

X-ray diffraction data was collected at 100 K at the MX2 and MX3 beamlines of the Australian Synchrotron. The XDS package was used for data processing and molecular replacement performed with Phaser. To solve the phase problem of PvAMA1-2700-864865 structural coordinates of PvAMA1, a human Fab and a nanobody and were used from PDB IDs 9DX6 and 7KJH. The phase problem of PvAMA1-2701 was solved using PvAMA1 and 2700 coordinates from PDB ID 9ZXJ. Strong translational non-crystallographic symmetry was detected for PvAMA1-2701 by Phaser with a Patterson peak 52.3 Å from the origin and a height of 48.7% relative to the origin. After tNCS correction the molecular replacement output had confident TFZ scores between 8.5-26.6 and LLG scores between 597-3744. Manual model building and structural refinement were conducted in alternating cycles by using COOT (version 0.9.8.96 EL) and Phenix (version 2.1-6048) [73,74].

### Data availability

Coordinates and structure factors of PvAMA1-2700-Fab 864865 and PvAMA1-2701 have been deposited in the Protein Data Bank under PDB IDs 9ZXJ and 44IE respectively.

## Acknowledgements

We would like to thank Roxanne Smith and the Bio21-WEHI Crystallisation Facility within Melbourne Protein Characterisation at The Bio21 Molecular Science and Biotechnology Institute, The University of Melbourne, as well as Geoffrey Kong and the Monash Macromolecular Crystallisation Platform (MMCP), for their assistance with setting up the crystallization screens. This research was undertaken using the MX2/MX3 beamline at the Australian Synchrotron, part of ANSTO, and made use of the Australian Cancer Research Foundation (ACRF) detector. We thank the MX2/MX3 beamline staff at the Australian Synchrotron for their assistance during data collection. The work was funded by The National Institutes of Health (R01AI143694) and Veterans Affairs Research Service (I01 BX001350). W-H.T. is supported by National Health and Medical Research Council of Australia (NHMRC) GNT2016908 and APP2001385. L-J.C was supported by NHMRC Investigator Grant GNT2041205. P.P is supported by NHMRC GNT2021267. JGB is supported by a NHMRC Investigator Grant (2033320). We thank the Australian Red Cross Lifeblood for the supply of human erythrocytes and serum. This work was made possible through the Victorian State Government Operational Infrastructure Support Program and the Australian Government NHMRC IRIISS.

---

**Figure S1.**
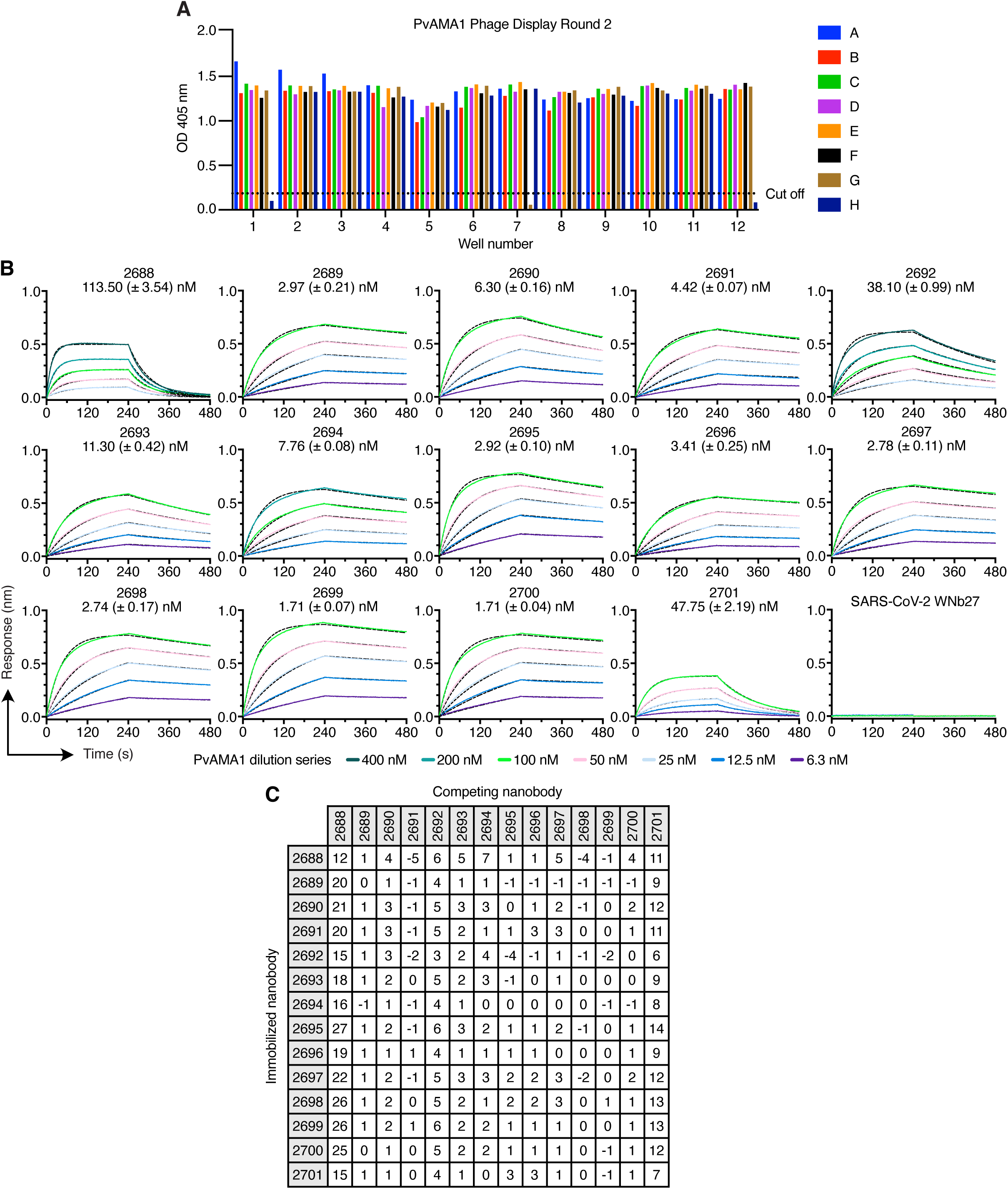
Phage ELISA screening of nanobodies and PvAMA1 nanobody affinities and epitope binning by BLI. **(A)** Monoclonal phage ELISA of individual nanobody-displaying phage clones following the second round of biopanning against PvAMA1. Bound phages were detected using a HRP-conjugated anti-M13 antibody, and absorbance was measured at 405 nm following ABTS development. The positive cut-off was defined as twice the absorbance of the negative controls, one non-relevant nanobody-displaying phage clone and one media only control (wells H1 and H12), indicated by the dashed line. Clones with absorbance values above the cut-off were considered positive binders. **(B)** BLI affinity curves using a two-fold serial dilution of PvAMA1 binding to immobilized nanobodies. Measurements were plotted (solid line) and fitted to a 1:1 binding model (black dashed line). *K*_D_ values represent means with standard deviations from two independent experiments (n=2) and a representative binding curve is shown from one experiment. **(C)** Epitope binning experiments of Sanger-selected nanobodies using BLI with PvAMA1 Palo Alto. Immobilized Sanger nanobodies are indicated on the left column and nanobodies pre-incubated with PvAMA1 Palo Alto at a 10:1 molar ratio are indicated on the top row. Binding of PvAMA1 pre-mixed with nanobody was calculated relative to PvAMA1 binding alone, which was assigned to 100%. The white boxes represent competing nanobodies.

**Figure S2.**
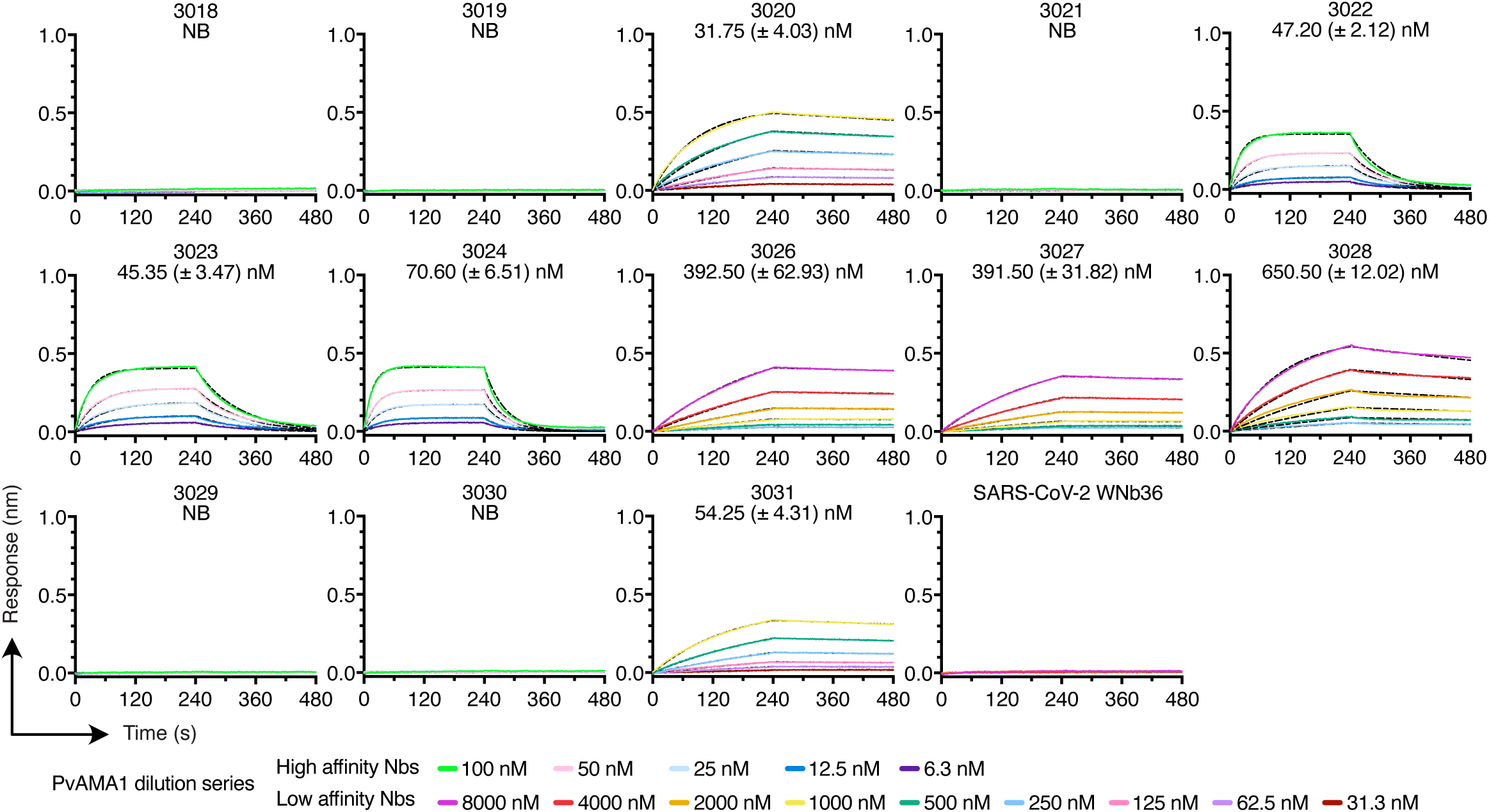
Affinities of AI-selected nanobodies to PvAMA1 by BLI. BLI affinity curves using a two-fold serial dilution of PvAMA1 binding to immobilized nanobodies. Measurements were plotted (solid line) and fitted to a 1:1 binding model (black dashed line). *K*_D_ values represent means with standard deviations from two independent experiments (n=2) and a representative binding curve is shown from one experiment. NB, no binding.

**Figure S3.**
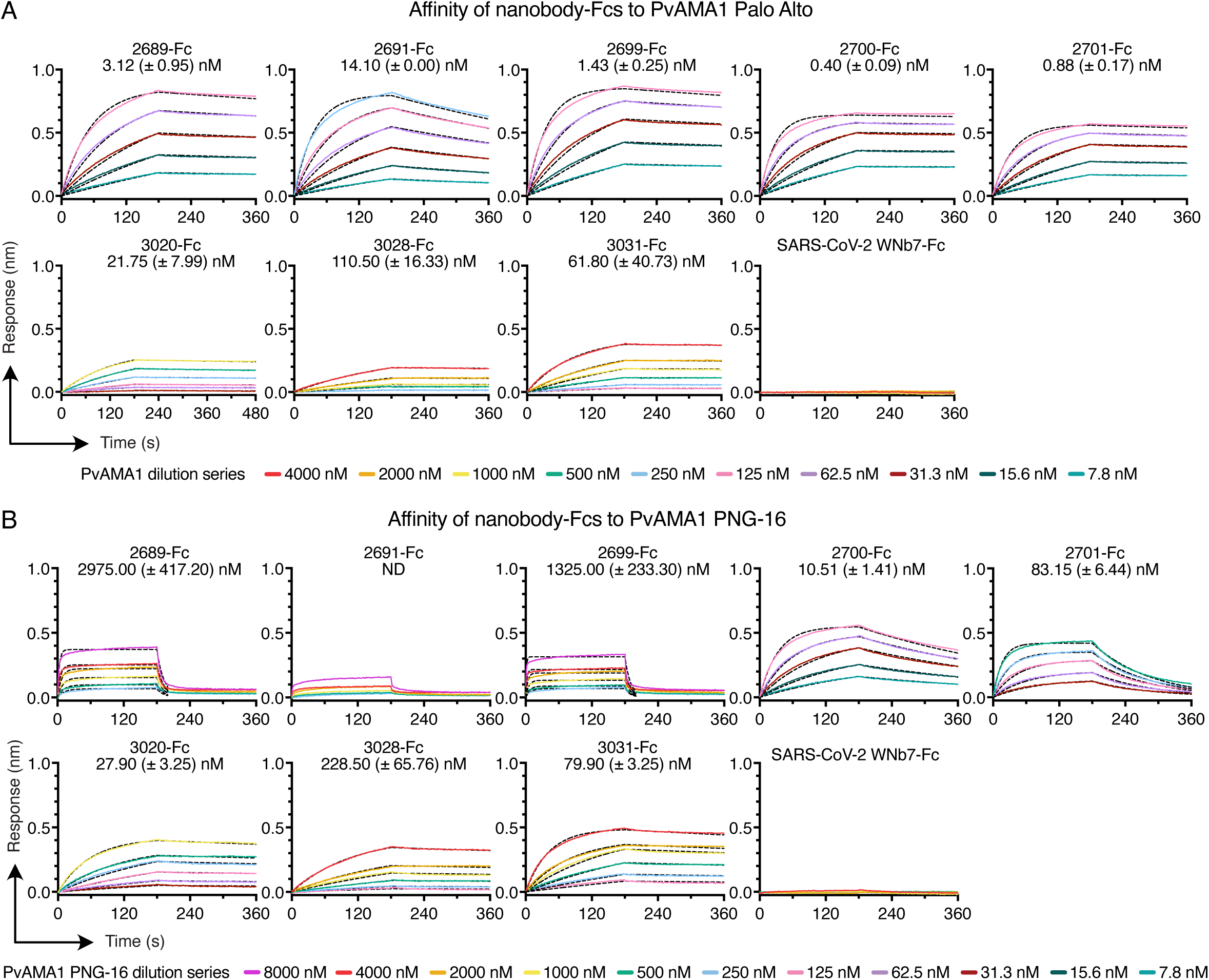
Affinities of nanobody-Fcs to PvAMA1 Palo Alto and PNG-16 by BLI. BLI affinity curves using a two-fold serial dilution of PvAMA1 Palo Alto or PNG-16 binding to immobilized nanobodies. Measurements were plotted (solid line) and fitted to a 1:1 binding model (black dashed line). *K*_D_ values represent means with standard deviations from two independent experiments (n=2) and a representative binding curve is shown from one experiment. ND, not determined due to poor fitting of the measured data to the binding model. All dissociation steps were performed for 180 s except for 3020-Fc that was performed at 300 s to obtain a dissociation measurement.

**Fig S4.**
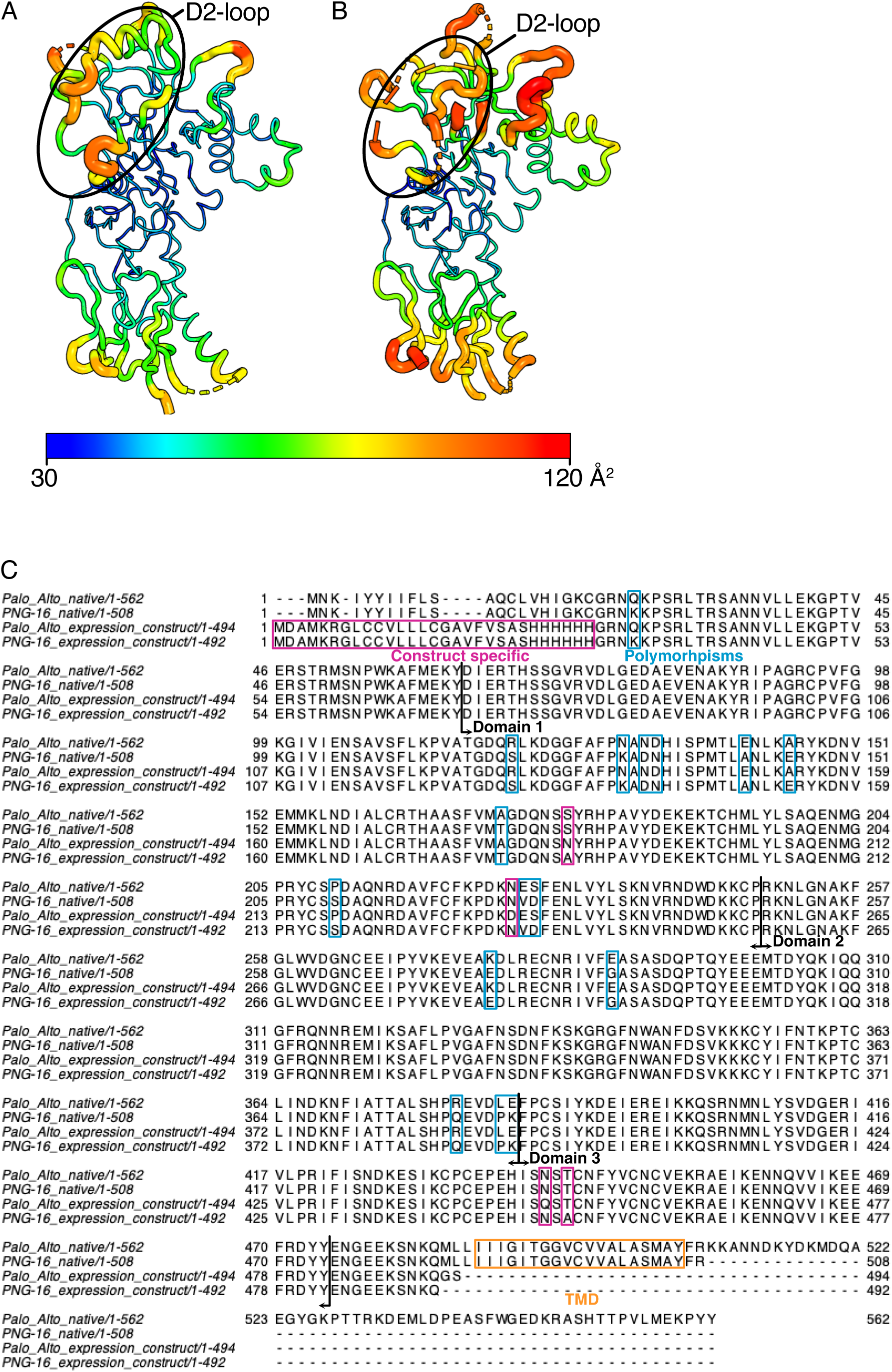
B-factor distribution in PvAMA1 of PvAMA1-2700-864865 (**A**) and of PvAMA1-2701 **(B)** structures shown as ‘putty’ representation. The C-alphaα B-factors are depicted on the structures with values from 30-120 Å^2^ from blue to red as indicated. The radius of the ribbon increases from low to high B-factor with the largest B-factors observed in the loop regions of D1 and the D2-loop. **(C)** Amino acid sequence alignments for PvAMA1 Palo Alto (ACB42433.1) and PvAMA1 PNG-16 (KC702426) native sequences alongside the expression construct specific sequences. Construct specific sequences and mutations are indicated in pink boxes. Polymorphisms are indicated in blue boxes. Transmembrane domain sequence is indicated in an orange box. Domain boundaries are indicated with black vertical lines and arrows.

**Figure S5.**
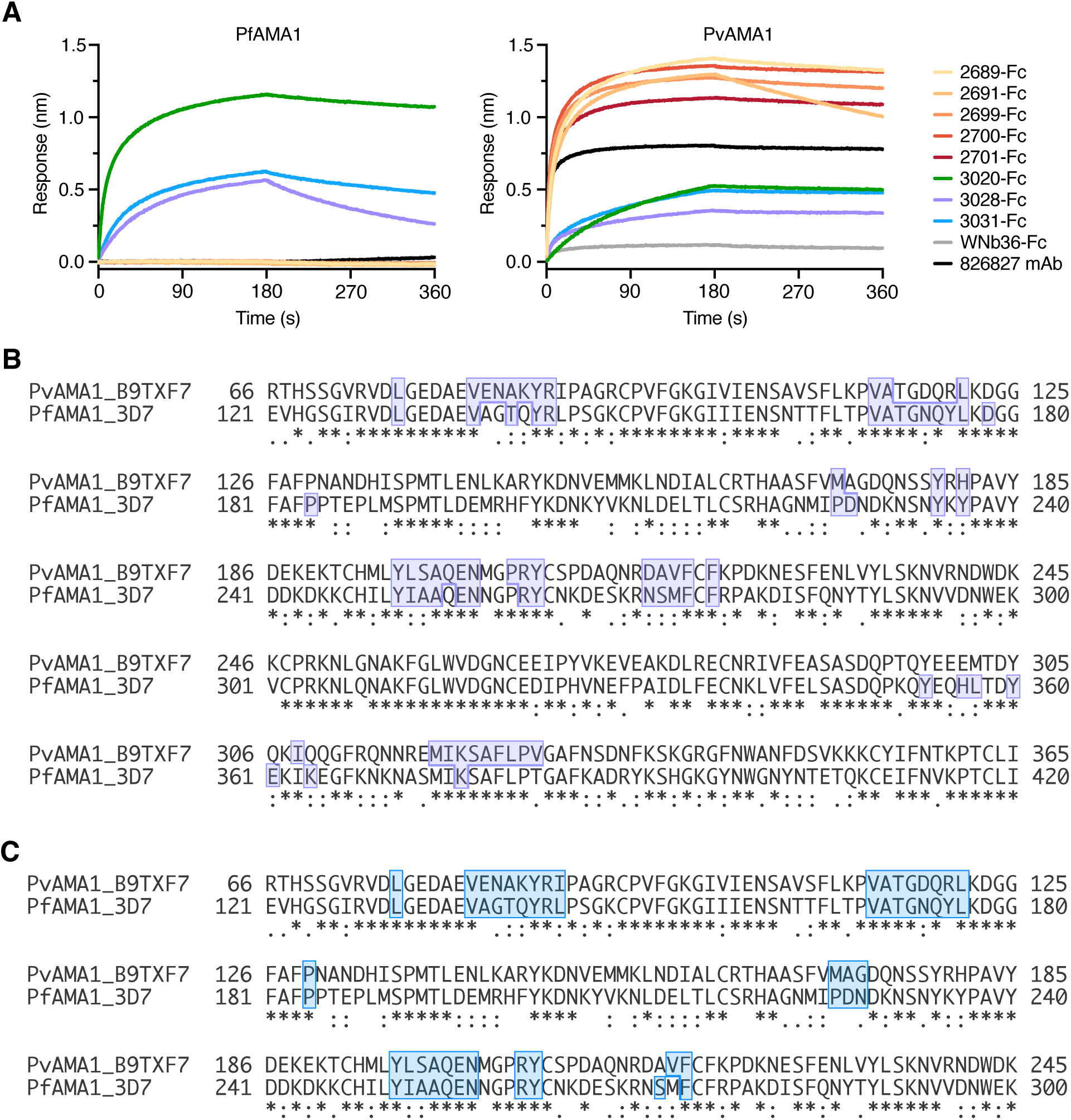
PvAMA1 nanobody-Fcs bind PfAMA1 and sequence conservation of predicted nanobody contact residues. **(A)** Binding analysis of PfAMA1 and PvAMA1 at a single concentration to nanobody-Fcs by BLI. Two independent experiments were performed with representative curves from one experiment shown. **(B)** Amino acid sequence alignment of PvAMA1 (from PDB 9DX6) and PfAMA1 (from 3D7 PDB 9EVO) with predicted contact residues for nanobody 3020 outlined with purple boxes. **(C)** Amino acid sequence alignment of PvAMA1 (from PDB 9DX6) and PfAMA1 (from 3D7 PDB 9EVO) with predicted contact residues for nanobody 3031 outlined with blue boxes.

**Data S1. Data relating to nanobody characterization performed in Figure 1 & 2 and S1 Fig & S2 Fig**. Tables of Sanger-selected and AI-selected NGS nanobody CDR3 sequences, full-length (FL) sequences, their corresponding counts per million (CPM) from NGS, enrichment from Sanger sequencing and determined binding kinetics and affinity data against PvAMA1 Palo Alto. Data are from two independent BLI experiments and shown are the mean ± SD values for affinity (*K*_D_), association rate (*k*_a_) and dissociation rate (*k*_d_). Chi-squared (X^2^) and R-squared (R^2^) are statistical values indicating quality of fit between experimental data and theoretical model. NB, no binding.

**Data S2. Data relating to methods for AI nanobody selection in Figure 2**. Tables of the top 100 most abundant (by counts per million, CPM) NGS nanobody CDR3 sequences after Round 1 and Round 2 of phage display. The GLG motifs are highlighted with orange text.

**Data S3. Data relating to methods for AI nanobody selection in Figure 2**. Table of AI-selected NGS nanobodies with CDR3 sequences, full-length (FL) sequences, their corresponding counts per million (CPM), their predicted binding site on PvAMA1, confidence metrics (ipTM and pTM) and interaction energies predicted using FoldX.

**Data S4. Data relating to parasite growth inhibition assays performed in Figure 3**. IC_50_ values calculated per biological replicate for PvAMA1 Sanger-selected and AI-selected nanobodies and nanobody-Fcs. Values given in both μg/mL and nM. Replicates are indicated as Rep 1, Rep 2 and Rep 3.

**Data S5. Data relating to nanobody characterization performed in Figure 4 & S3 Fig.** Binding kinetics and affinity data of strong nanobody inhibitors against PvAMA1 PNG-16 and nanobody-Fcs against PvAMA1 Palo Alto and PNG-16. Data are from two independent BLI experiments and shown are the mean ± SD values for affinity (*K*_D_), association rate (*k*_a_) and dissociation rate (*k*_d_). Chi-squared (X^2^) and R-squared (R^2^) are statistical values indicating quality of fit between experimental data and theoretical model. ND, not determined.

**Data S6. Data relating to parasite growth inhibition assays performed in Figure 6**. IC_50_ values calculated per biological replicate for PvAMA1 nanobodies and nanobody-Fcs when assessed in GIAs agasint *P. falciparum* strain 3D7. Values given in both μg/mL and nM. Replicates are indicated as Rep 1, Rep 2 and Rep 3.

**Data S7. Data relating to nanobody-Fc affinity measurements performed in Figure 6**. Binding kinetics and affinity data of nanobody-Fcs against PfAMA1. Data are from two independent BLI experiments and shown are the mean ± SD values for affinity (*K*_D_), association rate (*k*_a_) and dissociation rate (*k*_d_). Chi-squared (X^2^) and R-squared (R^2^) are statistical values indicating quality of fit between experimental data and theoretical model.

**Supporting Information**. Protein Data Bank validation report for 9ZXJ.

**Supporting Information**. Protein Data Bank validation report for 44IE.

**Table S1.** PISA generated interactions for PvAMA1 with 2700.

| PvAMA1 | Group | 2700 | Group | Distance (Å) |
|---|---|---|---|---|
| *Hydrogen bonds* |  |  |  |  |
| Glu 104 | OE2 | Ser 27 | OG | 2.9 |
| Met 203 | O | Arg 30 | NH1 | 3.8 |
| Glu 275 | O | Thr 57 | N | 3.2 |
| Glu 275 | N | Thr 57 | O | 2.7 |
| Glu 281 | OE1 | Asn 52 | ND2 | 3.1 |
| Glu 281 | OE1 | Ser 56 | OG | 2.8 |
| Arg 284 | NH2 | Asn 32 | OD1 | 2.8 |
| Glu 288 | OE2 | Asn 32 | ND2 | 3.7 |
| Glu 288 | OE1 | Leu 102 | N | 2.8 |
| Glu 288 | OE1 | Gly 103 | N | 2.9 |
| Tyr 298 | OH | Arg 100 | NH2 | 3.3 |
| Tyr 298 | OH | Glu 111 | OE1 | 2.7 |
| *Salt bridges* |  |  |  |  |
| Glu 104 | OE1 | Arg 30 | NE | 3.2 |
| Glu 104 | OE2 | Arg 30 | NE | 3.1 |
| Glu 104 | OE1 | Arg 30 | NH2 | 2.9 |
| Glu 201 | OE2 | Arg 100 | NE | 2.7 |
| Glu 201 | OE1 | Arg 100 | NE | 3.8 |
| Glu 201 | OE2 | Arg 100 | NH2 | 2.8 |
| Asp 226 | OD1 | Arg 54 | NH2 | 3.4 |
| Other 2700 interfacing residues (PvAMA1) |  |  |  |  |
| Arg 73 | Val 102 | Ile 103 | Asn 105 | Cys 208 |
| Ser 209 | Pro 210 | Cys 220 | Phe 221 | Lys 222 |
| Pro 223 | Asp 224 | Glu 227 | Lys 272 | Glu 273 |
| Val 274 | Ala 276 | Lys 277 | Arg 280 | Ile 285 |
| Ala 289 | Ser 292 | Gln 294 | Thr 296 | Phe 342 |
| Ile 356 |  |  |  |  |
| Other PvAMA1 interfacing residues (2700) |  |  |  |  |
| Gln 1 | Phe 29 | Asp 53 | Gly 55 | Tyr 58 |
| Tyr 59 | Pro 99 | Gly 101 | Tyr 104 | Met 106 |
| Thr 107 |  |  |  |  |

The distance measurements are based on molecules A and G.
Interacting and interfacing residues between PvAMA1, antibody Fab 864865 and nanobody 2700 were determined using PISA.
Some residue side chains are unresolved and the interactions listed here may not be complete.

**Table S2.** PISA generated interactions for PvAMA1 with and 2701.

| PvAMA1 | Group | 2701 | Group | Distance (Å) |
|---|---|---|---|---|
| *Hydrogen bonds* |  |  |  |  |
| Glu 104 | OE2 | Tyr 32 | OH | 2.7 |
| Met 203 | O | Arg 101 | NH2 | 3.2 |
| Pro 205 | O | Tyr 115 | OH | 3.2 |
| Lys 222 | NZ | Pro 100 | O | 2.8 |
| Lys 222 | NZ | Ser 31 | OG | 2.8 |
| Glu 275 | N | Thr 58 | O | 2.8 |
| Glu 275 | O | Thr 58 | N | 2.9 |
| Glu 275 | OE1 | Thr 58 | OG1 | 3.8 |
| Lys 277 | N | Ala 56 | O | 3.1 |
| Glu 288 | OE1 | Gly 104 | N | 3.2 |
| Glu 288 | OE1 | Leu 103 | N | 3.6 |
| *Salt bridges* |  |  |  |  |
| Glu 201 | OE2 | Arg 101 | NE | 3.0 |
| Glu 273 | OE1 | Lys 65 | NZ | 2.8 |
| Glu 281 | OE1 | Arg 52 | NH2 | 3.3 |
| Glu 281 | OE2 | Arg 52 | NE | 3.4 |
| Glu 288 | OE1 | Arg 52 | NH2 | 2.9 |
| Glu 288 | OE2 | Arg 52 | NH1 | 3.0 |
| Other 2701 interfacing residues (PvAMA1) |  |  |  |  |
| Arg 73 | Val 102 | Asn 105 | Gly 204 | Pro 223 |
| Asp 224 | Tyr 270 | Lys 272 | Val 274 | Ala 276 |
| Arg 284 | Ile 285 | Ala 289 | Ser 290 | Ala 291 |
| Ser 292 | Gln 294 | Thr 296 | Gly 341 | Phe 342 |
| Ile 356 |  |  |  |  |
| Other PvAMA1 interfacing residues (2701) |  |  |  |  |
| Arg 27 | Phe 29 | Ser 30 | Ser 54 | Gly 55 |
| Ser 57 | Tyr 59 | Tyr 60 | Asp 62 | Gly 102 |
| Tyr 105 | Ile 107 | Arg 108 | Asp 109 | Asp 144 |

The distance measurements are based on molecules A and G.
Interacting and interfacing residues between PvAMA1 and nanobody 2701 were determined using PISA.
Some residue side chains are unresolved and the interactions listed here may not be complete.

